# Inference of heterogeneous effects in single-cell genetic perturbation screens

**DOI:** 10.1101/2024.05.28.596224

**Authors:** Zichu Fu, Lin Hou

## Abstract

Recent single-cell CRISPR screening experiments have combined the advances of genetic editing and single-cell technologies, leading to transcriptome-scale readouts of responses to perturbations at single-cell resolution. An outstanding question is how to efficiently identify heterogeneous causal effects of perturbations using these technologies. Here we present scCAPE, a tool designed to facilitate causal analysis of heterogeneous perturbation effects at the single-cell level. scCAPE disentangles perturbation effects from the inherent cell-state variations and provides nonparametric inferences of perturbation effects at single-cell resolution, permitting a range of downstream tasks including perturbation effect analysis, genetic interaction analysis, perturbation clustering and prioritizing. We benchmarked scCAPE through simulation studies and real datasets to evaluate its performance in characterizing latent confounding factors and accuracy in estimating heterogeneous perturbation effects. The application of scCAPE identified novel heterogeneous genetic interactions among erythroid differentiation drivers. For example, our analysis pinpointed the role of the synergistic interaction between CBL and CNN1 in the S phase.

## Introduction

Experimentally perturbing a gene and quantifying the phenotypic responses is an important method for studying gene functions. By selectively introducing CRISPR edits to many genes or genomic sites at the same time, the CRISPR screening experiments have facilitated pinpointing many key genes involved in cellular differentiation, immune responses, disease mechanisms and more^1^. Recent single-cell CRISPR screening technologies like Perturb-seq^2,3^, CRISP-seq^4^ and CROP-seq^5,6^ enhanced the capabilities of perturbation screening by combining CRISPR screening with single-cell RNA sequencing (scRNA-seq). Through the association of guide RNAs (gRNAs) with individual cell transcriptomes, these methods expand the low-content readouts like cell growth or marker expression^7–10^ to rich molecular profiles at the resolution of single cells, providing more elaborated information about the gene functions and regulations.

To quantify perturbation effects, the transcriptional profiles of cells with and without perturbations are compared. Before conducting this comparison, we need to make several important decisions. First, what resolution should be employed for the comparison? Most current approaches focus on estimating average perturbation effects at the pseudo-bulk level; however, it is intriguing to investigate whether the perturbation effect can be assessed at the single-cell level. Pseudo-bulk approaches implicitly assume homogeneity of perturbation effects across cells. While pooling cells enhances statistical power in average effect approaches, it does so at the expense of disregarding the potential heterogeneous landscape of perturbation effects which can offer valuable insights into regulatory mechanisms. For instance, knocking out UBASH3A shows a notable increase in proliferation in TCR (T cell receptor)-stimulated T cells but not in unstimulated cells, indicating the gene’s regulatory effect on proliferation is induced by TCR-signaling^11^. At first glance, it may seem straightforward to estimate heterogeneous perturbation effects by analyzing subclusters of cells separately^4,11^. However, in doing so, one needs to make arbitrary decisions on segmenting cell populations, which easily suffers from data snooping bias^12,13^ and sacrifices the power of single-cell data. Thus, a framework that adaptively partitions cells and estimates perturbation effects at single-cell resolution is necessary to probe the intricacies of gene regulation.

Second, what phenotype should we compare, given the readout of the entire transcriptome? An obvious choice could be the expression of every gene^3,14,15^. However, conducting individual-gene level analysis brings the burden of multiple testing, and the statistical power is limited due to the high level of noise in scRNA-seq data and the relatively small number of cells per perturbation in typical experiments. Recently, methods such as MUSIC and GSFA have explored perturbation effects on gene modules^16,17^, motivated by the observation that genetic perturbations tend to influence the expression of multiple related genes simultaneously, rather than exclusively targeting individual genes^18^. Compared with individual-gene level analysis, examining perturbation effects on these gene modules, which represent specific pathways or functional clusters, can enhance statistical powers by aggregating the effects across genes and offer a more explicit biological understanding of the roles of perturbed genes.

Third, how do we characterize and correct for confounding factors? In single-cell CRISPR screening data, a significant proportion of transcriptomic variations are attributed to sources other than the perturbation^3,19^. Sources such as cell types and experimental conditions can be easily identified and handled, while many other sources can be relatively subtle and challenging to detect and model explicitly. For example, in the studies involving differentiating cell lines such as K562 cells^3,20^ and Lund mesencephalic cells (LUHMES)^21^, differentiation states and cell cycle stages can both act as confounding factors. Ignoring these inherent cell state variations can lead to false discovery of associations between perturbations and gene expression or reduce the power to identify true perturbation effects. Therefore, it is crucial to disentangle perturbation effects from confounding factors in the transcriptome and adjust for these confounders when estimating perturbation effects.

Here, we present scCAPE, a framework that enables Causal Analysis of heterogeneous Perturbation Effects in single-cell perturbation screening data. Our method efficiently disentangles perturbation effects from the inherent cell state variations and estimates perturbation effects at single-cell resolution. We benchmark scCAPE using simulations and real datasets and demonstrate its applications in identifying genetic interactions, as well as clustering and prioritizing perturbation effects.

## Results

### scCAPE method overview

scCAPE is designed to estimate heterogeneous causal effects in genetic perturbation data. Its input comprises normalized transcriptome profiles and perturbation labels from single-cell CRISPR screening experiments.

A key component of scCAPE is an autoencoder with adversarial training to disentangle perturbation effects from confounding factors in the transcriptome profiles. The goal is to identify transcriptomic changes that are the causal consequences of introducing a particular perturbation and would otherwise not be observed. Through disentangling, we seek a latent embedding space to characterize intrinsic and/or extrinsic covariates of each cell (referred to as a basal space), which represents the ‘pre-perturbed’ cell status and contains no information about perturbations (see **Figure 1**). Based on the basal space, subsequent causal components of scCAPE can statistically match perturbed and unperturbed cells to make unbiased inferences of perturbation effects. To achieve disentanglement, scCAPE trains a discriminator to predict perturbation labels, which competes against the encoder that thwarts the discriminator’s prediction ability (**Methods** and **Supplementary Note**).

**Figure 1.**
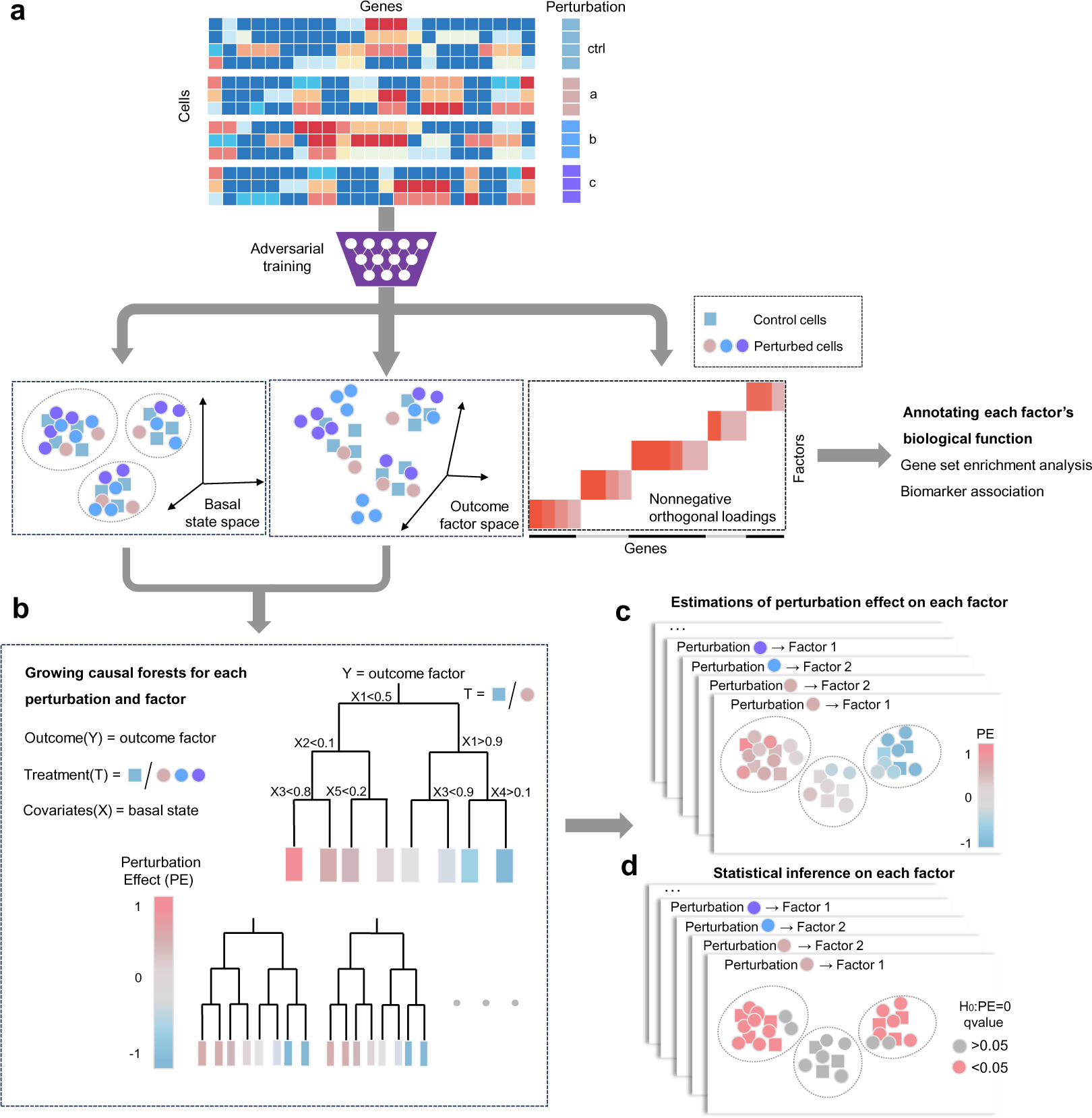
Overview of scCAPE workflow. scCAPE employs a two-step modeling to infer the heterogeneous perturbation effects. Step 1: (a) Disentangling the perturbation effects from inherent cell-state variations through an autoencoder framework with adversarial training. The neural networks output the basal state and the outcome factors of each cell. The biological functions of each factor can be inferred based on the nonnegative orthogonal gene loadings. Step 2: (b) Growing causal forests for each pair of perturbation and factor. (c) Through iteratively splitting the basal space, the causal trees provide the estimations of perturbation effects on each factor at single-cell resolution. (d) The asymptotic variances of the estimators can then be used to conduct hypothesis tests.

Then, the decoder module combines the basal space and the perturbation labels to learn outcome factors, which are subsequently used to reconstruct the transcriptome profiles. These outcome factors summarize the expression of multiple related genes and align well with known biological functions. We impose non-negative and orthogonal constraints on the factor loading matrix for better interpretation, and biological functions are annotated through biomarker associations and gene set enrichment analysis (**Methods**).

Having obtained the embeddings disentangled from the neural networks, we estimate the perturbation effects for each perturbation-factor pair by growing causal forests, as illustrated in **Figure 1**. By iteratively splitting the basal space into subspaces that maximize perturbation effect heterogeneity, we obtain estimations and asymptotic variances of perturbation effects for each cell. Subsequently, statistical tests are conducted to show if the perturbation effects are significant (**Methods**).

Finally, with the perturbation effects at single-cell resolution, scCAPE can quantitively characterize the roles of perturbed genes at any subpopulation of cells from the following perspectives (**Figure 7** and **Methods**): quantifying the relationships between different perturbations and prioritizing the perturbations based on overall perturbation effects, or in a functional factor-specific way. Furthermore, scCAPE is extended to facilitate the analysis of genetic interactions in combinatorial perturbation datasets (**Figure 6** and **Methods**).

We demonstrate the power of scCAPE by benchmarking it on simulated and real datasets. Using single-cell CRISPR data from human CD8+ T cells and K562 cells, we dissected the relationship between perturbation responses and T cell receptor stimulation and revealed heterogeneous perturbation landscapes of cell cycle regulators. Another application of scCAPE on a combinatorial perturbation dataset identified diverse genetic interactions across different pathways and uncovered strengthened synergistic effects between erythroid drivers in the S stage.

### Evaluating the reconstruction and disentangling performance

We applied scCAPE to a dataset of human peripheral blood mononuclear cells (PBMCs) stimulated with interferon (IFN-β)^22^ (**Figure 2a**), consisting of seven cell types, 8886 stimulated cells and 8007 control cells. We scrutinized the reconstruction errors of the autoencoder framework in scCAPE which introduces orthogonal constraints and adversarial training. This was measured by squared Pearson correlation (*R*^2^) between the reconstruction and the original data (**Figure 2c**). The reconstructed profiles align well with the real data, with average *R*^2^ of 0.939 and 0.933 for all genes and the top 100 differentially expressed genes across cell types. For example, the average reconstruction *R*^2^ is 0.976 in the stimulated F-Mono cells (**Figure 2b**).

**Figure 2.**
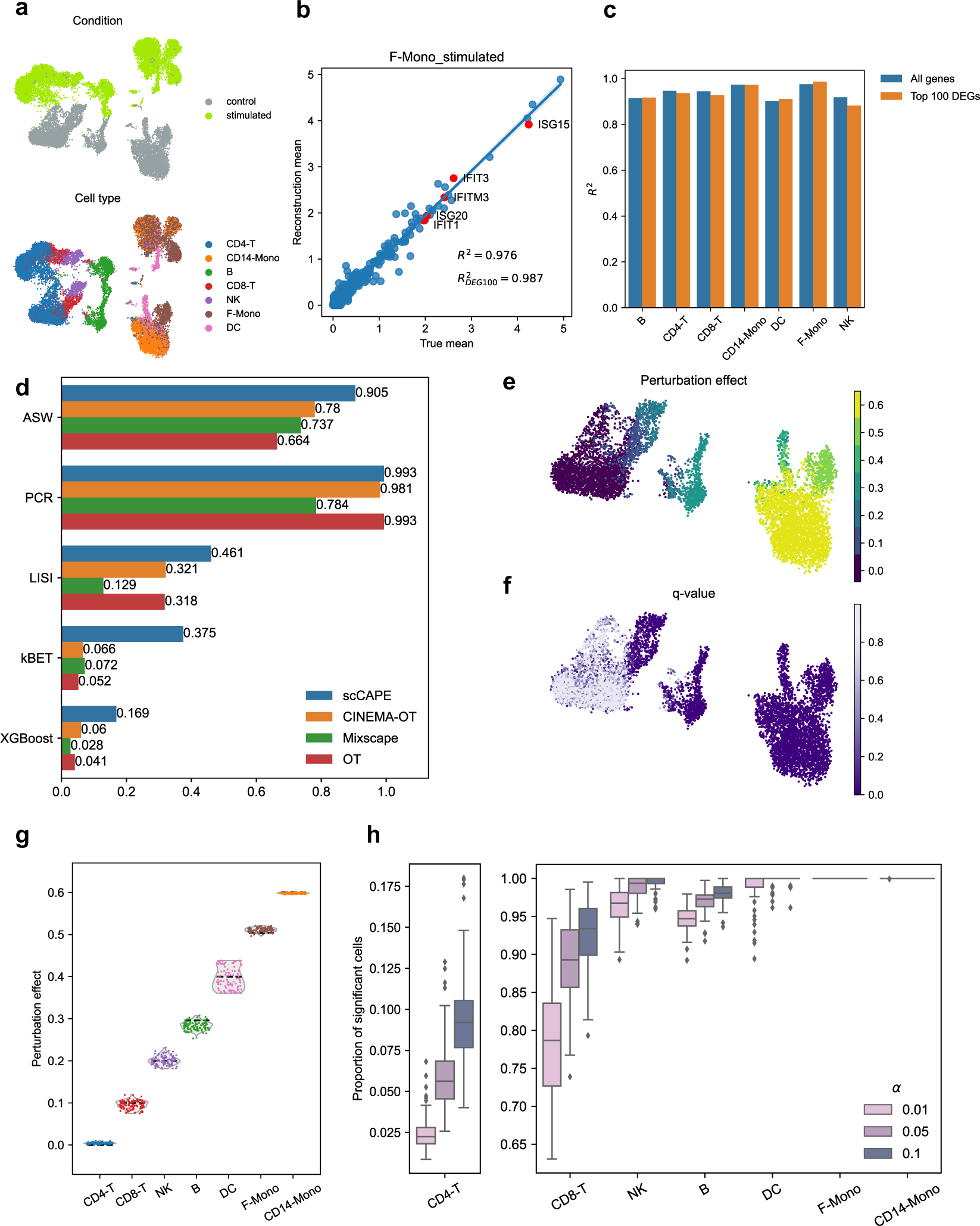
The validation of scCAPE on the PBMC dataset. (a) UMAP visualizations^53^ of the dataset, colored by conditions and cell types. (b) Mean gene expression of scCAPE reconstructed and real in stimulated F-Mono cells. The red points represent the top five upregulated DEGs. *R*^2^ denotes squared Pearson correlation between real and reconstructed values. (c) Comparison of *R*^2^ values for mean gene expression between real and reconstructed for the seven cell types. (d) The mixing performance of perturbed and control cells in the embedding of scCAPE, CINEMA-OT, OT and Mixscape. (e) UMAP visualization of cells in the simulation dataset, colored by perturbation effects estimated by scCAPE. (f) UMAP visualization of cells in the simulation dataset, colored by q-values inferred by scCAPE. (g) Violin plots of perturbation effects within seven cell types estimated by scCAPE. For each violin, *n* = 100 estimates generated from 100 rounds of simulation under the setup. The dotted lines represent the true perturbation effect in each cell type. (h) Box plots of the proportion of significant cells within seven cell types inferred by scCAPE. The different colors represent different values of significance level (*α*) varying from 0.01 to 0.1. For each box, *n* = 100 results generated from 100 rounds of simulation under the setup.

Next, we evaluated the disentangling effect of the basal state. Ideally, the basal space is a low-dimensional embedding that bears the inherent cell status prior to any perturbation. We note that previous methods^23,24^ achieve similar goals by optimal matching between perturbed cells and control cells. scCAPE directly implements adversarial training to learn a perturbation-invariant basal state that does not carry information of the perturbation labels. Our approach performs well in preserving the cell type information (**Supplementary Figure 2**), and is advantageous in mixing cells regarding perturbation labels (**Figure 2d**). More importantly, the perturbation-invariant embedding enables causal inference of the transcriptomic responses to perturbations.

### Evaluating the performance of perturbation effect inference

We performed simulation studies based on the PBMC dataset to demonstrate that scCAPE not only provides unbiased estimations of perturbation effects but also enables effective hypothesis testing. Specifically, we divided the control cells in the PBMC dataset randomly into two equal subsets. One subset served as the control group, while for the other subset, we introduced perturbation effects at the outcome factor level. To introduce heterogeneity in perturbation effects, their magnitudes were varied across the seven cell types, ranging from 0 to 0.6 (**Methods**). Subsequently, we estimated the perturbation effects at the single-cell level (**Figure 2e**) and averaged them across the seven cell types separately. Hypothesis testing was also conducted to assess the significance of the estimated effects (**Figure 2f**). Our results demonstrate that scCAPE accurately estimates the heterogeneous perturbation effects (**Figure 2g**) and effectively identifies cells with non-zero perturbation effects (**Figure 2h**). For instance, in CD4-T cells where no perturbation effects were introduced, the estimates are around zero and the tests have low rejection rates. While in CD8-T cells subjected to minor perturbation effects, scCAPE successfully captures the small effects and possesses sufficient power to reject the null hypothesis. In addition, we assessed the calibration of scCAPE’s estimates and inference results by randomly permuting the perturbation labels (**Supplementary Figures 4 and 5**). At the significance level of 0.05, the proportion of falsely detected cells was less than 0.01.

In sum, the autoencoder framework of scCAPE successfully reconstructs the data and identifies a basal space to characterize cellular heterogeneity prior to perturbations that is essential for causal inference of perturbation effects. scCAPE estimates perturbation effects in an unbiased manner and demonstrates good statistical power and limited type-I error in hypothesis testing of the perturbation effects.

### scCAPE reveals heterogeneous perturbation landscapes of T cell regulators

We applied scCAPE to a CROP-seq dataset that investigated the mechanisms of stimulation response in primary human CD8+ T cells^11^, where fourteen T cell regulators and six well-established immune checkpoint genes were targeted using CROP-seq in both T cell receptor (TCR) stimulated and unstimulated T cells (**Figure 3c**).

**Figure 3.**
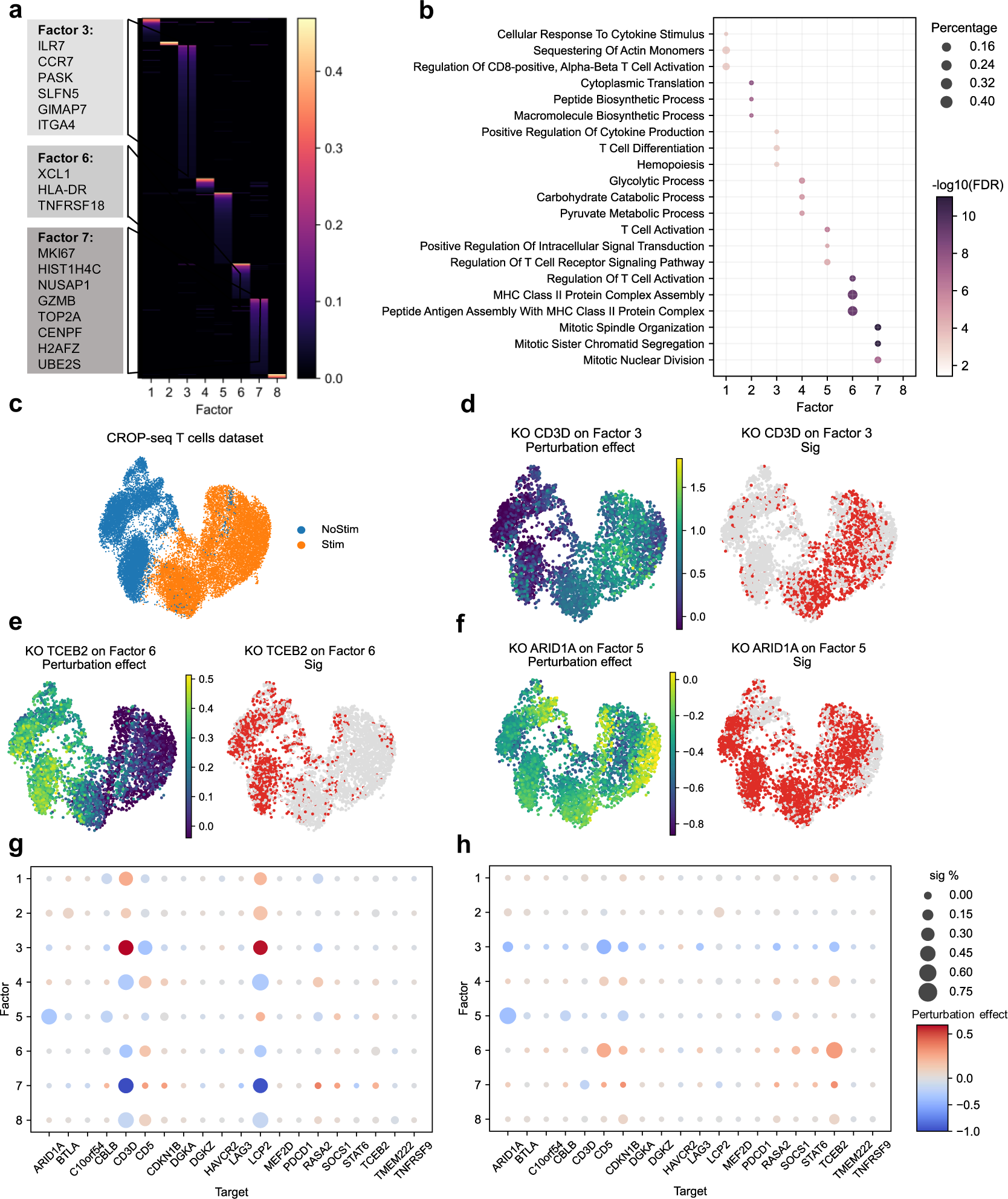
scCAPE reveals heterogeneous perturbation effects in the CROP-seq T cells dataset. (a) The nonnegative and orthogonal loading matrix annotated with some markers genes. (b) The Gene Ontology (GO) enrichment analysis^25^ of gene sets selected from each factor. The size of the dot represents the percentage of selected genes in that GO term. The color of the dot shows the −log_10_(adjusted p-value). (c) UMAP visualization of stimulated cells (Stim) and unstimulated cells (NoStim).(d-f) The different patterns of heterogeneous perturbation effects. ‘Sig’ indicates whether the perturbation effect is significant (red, q-value<0.05) or not (grey, q-value>0.05). These results were further validated by comparing the values of factors between perturbed and control cells in stimulated and unstimulated states separately (**Supplementary Figure 8**). (g, h) The perturbation effect of each perturbation on each factor, aggregated in stimulated cells (g) and unstimulated cells (h), respectively. The color represents the average perturbation effects and the size indicates the percentage of cells with significant perturbation effects (q-value<0.05). In stimulated cells, many associations between perturbations and factors that are significant in large proportions of cells, while the perturbation effects in unstimulated cells are relatively subtle.

Eight outcome factors were selected and annotated with specific biological functions (**Figure 3a**-**b**, and **Supplementary Figure 7**). For example, factor 3 has large weights for T cell resting markers such as ILR7 and CCR7. Factor 6 is heavily weighted with T cell activation markers XCL1 and TNFRSF18. Factor 7 encompasses T cell proliferation markers, including MKI67, TOP2A, and CENPF, and exhibits enrichment for Gene Ontology (GO) terms^25^ cell cycle and division.

For each factor, scCAPE estimated the perturbation effect for each cell, revealing a heterogeneous landscape of perturbation effects. For instance, the knockout of CD3D exhibits positive effects on factor 3 exclusively in stimulated cells (**Figure 3d**), whereas the knockout of TCEB2 exerts its effects solely in unstimulated cells (**Figure 3e**), indicating their regulatory effects are dependent on TCR signaling. Moreover, we discovered that ARID1A, a target poorly characterized in the original study^11^, functions as a positive regulator of T cell activation in both stimulated and unstimulated cells (**Figure 3f**). Indeed, ARID1A has gained recognition as a chromatin remodeler and tumor suppressor across a broad spectrum of cancers^26,27^, with its mutation associated with poor immunotherapy response and shortened host survival^28,29^. We summarized the single-cell resolution results for these two conditions respectively (**Figure 3g-h**). In comparison with the individual gene-level analyses conducted in the original study^11^, the clear interpretations of the factors in scCAPE allow for a direct understanding of the perturbation effects. For example, gRNAs targeting CD3D and LCP2 both increase the expression of T cell resting marker genes while significantly decreasing the expression of T cell activation and proliferation marker genes in most stimulated cells. Conversely, the knockout of CD5 produces the opposite effects.

In addition to exploring the heterogeneous effects between the two conditions, the perturbation map generated by scCAPE enables us to examine at a finer resolution, thereby revealing subpopulations of cells with distinct perturbation effects. For instance, targeting LCP2 significantly decreases the level of factor 7 (T cell proliferation) in stimulated cells, which was reported in previous studies^11,17^. Furthermore, scCAPE uncovers that this effect is distributed across a broad range, with similar effect sizes observed within subpopulations of cells (**Figure 4a-b**). By employing the Leiden algorithm^30^ in the basal space, we identified four subgroups of stimulated cells (**Figure 4c**), with variations in perturbation effects aligning closely with these cell subpopulations. Notably, although cluster 1 cells are stimulated, their transcriptome resembles that of unstimulated cells. Similar to unstimulated cells, targeting LCP2 exhibits no significant effect for the majority of cells in this cluster (**Figure 4d-e**). We observed that variations in perturbation effects among the four clusters can primarily be attributed to differences in stimulation intensity, as evidenced by correlation analysis between perturbation effects and the expression of markers indicative of the unstimulated state (CCR7 and ILR7) and stimulated state (GZMB, MKI67)^11^ (**Figure 4f**). We also found similar associations between the extent of perturbation effects and the intensity of stimulation for other target genes (**Supplementary Figure 10**).

**Figure 4.**
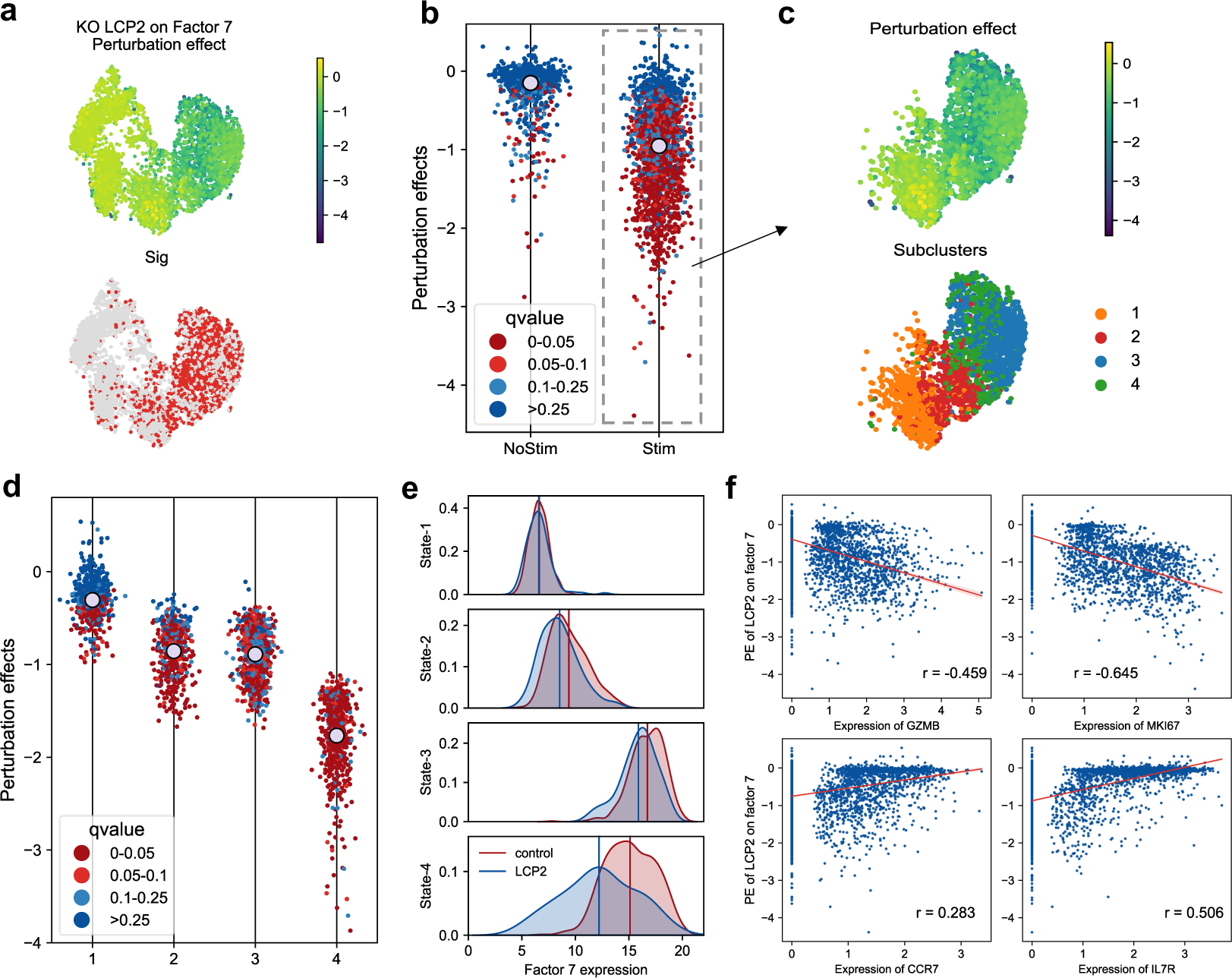
scCAPE characterizes subpopulations that respond differently to perturbations. (a) The perturbation effect and significance of the knockout (KO) of LCP2 on factor 7 (cell proliferation). (b) The perturbation effect of KO LCP2 on factor 7, aggregated on stimulated (Stim) and unstimulated (NoStim) cells. The color of each cell represents the q-value. The grey dot shows the average perturbation effects on each condition. The stimulated cells exhibit large ranges of perturbation effects, suggesting non-uniform responses to the perturbation. (c) UMAP visualization of stimulated cells, colored by the perturbation effects and the subclusters labels. (d, e) We observed that although cells in subcluster 1 are stimulated, the perturbation effect is not significant (d). This finding is also validated by comparing the expression of factor 7 in different subclusters (e). (f) We performed linear regression of the perturbation effect on the expression of four marker genes (GZMB, MKI67, CCR7 and IL7R). Each dot represents a control cell. *r* denotes the correlation coefficient.

Collectively, scCAPE can explore the heterogeneity of perturbation effects at various levels of granularity without the necessity of predefining cell subgroups, thus unveiling the intricate molecular effects of genetic perturbations in single-cell CRISPR screens.

### scCAPE reveals heterogeneous regulatory effects of cell-cycle regulators

We next applied scCAPE to a Perturb-seq dataset that investigated the mechanism of 13 potential cell cycle regulators in K562 cells^3^, wherein the original study revealed different cellular states can confound the study of perturbation effects^3^. We set out to investigate the perturbation effects of these regulators on signature gene sets^31^ of different cell-cycle phases.

scCAPE selected 15 factors and conducted gene set enrichment analysis to annotate their functions (**Supplementary Figure 12**). Then the factor level perturbation effect estimates were transformed to the individual gene level using factor loadings (**Methods**) and subsequently averaged on the signature gene sets of each cell-cycle phase. We employed the Leiden algorithm^30^ on our basal embedding to categorize cells into four states. The differential expression and gene set enrichment analysis revealed their biological distinctions (**Figure 5c**). For instance, cell states 2 and 3 are associated with erythroid differentiation, wherein cells in state 2 exhibit high expression of hemoglobin genes, while those in state 3 exhibit high expression of ribosomal protein genes which facilitates sufficient ribosome production for globin translation during early erythropoiesis^32^. Cells in state 4 upregulate pathways related to endoplasmic reticulum (ER) stress.

**Figure 5.**
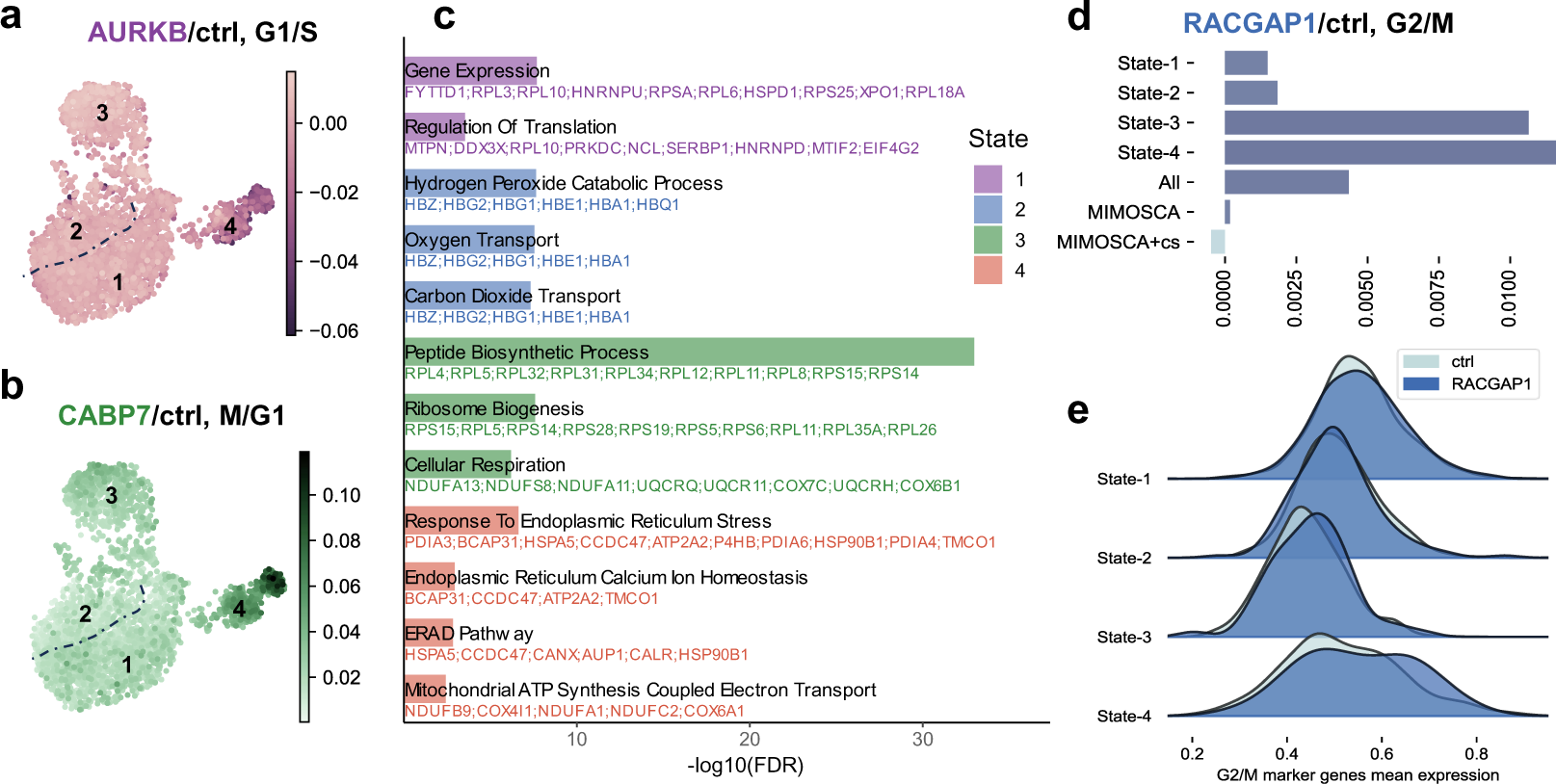
scCAPE reveals heterogeneous perturbation effects in the Perturb-seq K562 cells dataset. (a) UMAP visualization of the perturbation effect of KO AURKB on G1/S signature gene sets, where the numbers represent the four cell states. (b) Similar plot to (a), displaying the perturbation effect of KO CABP7 on M/G1 signature gene sets. (c) Gene ontology (GO) enrichment results of the top 50 differentially expressed genes in each cell state. (d) The perturbation effect of KO RACGAP1 on G2/M signature gene sets aggregated across 4 states and all cells, with the average perturbation effect estimation obtained from MIMOSCA. ‘MIMOSCA+cs’ denotes MIMOSCA fitting with cell-state covariates. (e) Density plots illustrating the average expression of G2/M signature genes in RACGAP1-perturbed cells and control cells aggregated across four cell states.

In comparison to the average perturbation effects estimated by the original study, the single-cell resolution estimations provided by scCAPE allowed for a more in-depth exploration of perturbation effects. For instance, on average, the knockout of AURKB was associated with a reduction in G1/S signatures. Our analysis further revealed that this decrease occurred specifically in state 4 (**Figure 5a**). Similarly, the knockout of CABP7 was linked to an increase in M/G1 signatures, with state 4 exhibiting a more pronounced increase (**Figure 5b**). We observed that perturbation effects appeared to be more prominent in state 4, suggesting that perturbations have a greater impact in an ER stressful environment. Given that ER stress and the associated unfolded protein response can trigger cell death, conditions leading to increased protein misfolding or decreased cellular capacity to manage these proteins in the ER may result in more severe perturbation impacts and cellular dysfunction^33^.

We noticed that the previous methods^3,15^ can also correct for cell status variations when estimating perturbation effects by fitting the linear regression with confounders as covariates. However, in these methods, the cell status must be predetermined, and the estimated effect is averaged over all cells. In contrast, scCAPE is data-adaptive, and the perturbation effect is estimated in single-cell resolution through a nonparametric approach. These advantages of scCAPE result in more precise estimations of perturbation effects. For instance, scCAPE revealed that the knockout of RACGAP1 was associated with an increase in G2/M signatures across all cell states (**Figure 5d-e**). Consistently, the knockout of RACGAP1 has been implicated in inducing G2/M phase arrest^34,35^. However, MIMOSCA failed to identify this association and even reversed the trend when cell-state covariates were included, indicating that inadequate modeling of confounding factors and neglecting the heterogeneity in perturbation effects can obscure genuine perturbation effects.

### scCAPE infers genetic interactions in high-MOI screens

In CRISPR screens with high multiplicity of infection (MOI), where one cell can be targeted by more than one gRNA, the inference framework of scCAPE can be extended to test for genetic interactions (GIs). To elaborate, we grow multi-armed causal forests to jointly infer the effect of one-gene perturbations and two-gene perturbations on each factor, and devise statistical tests for genetic interactions (**Figure 6a-b**). Here we focus on the analysis of four types of GIs^3,20^: dominant, synergistic, buffering, and epistasis (**Methods**). It is worth mentioning that the power of scCAPE is not limited to the four types of GIs, as it can easily accommodate other types of GIs of interest.

**Figure 6.**
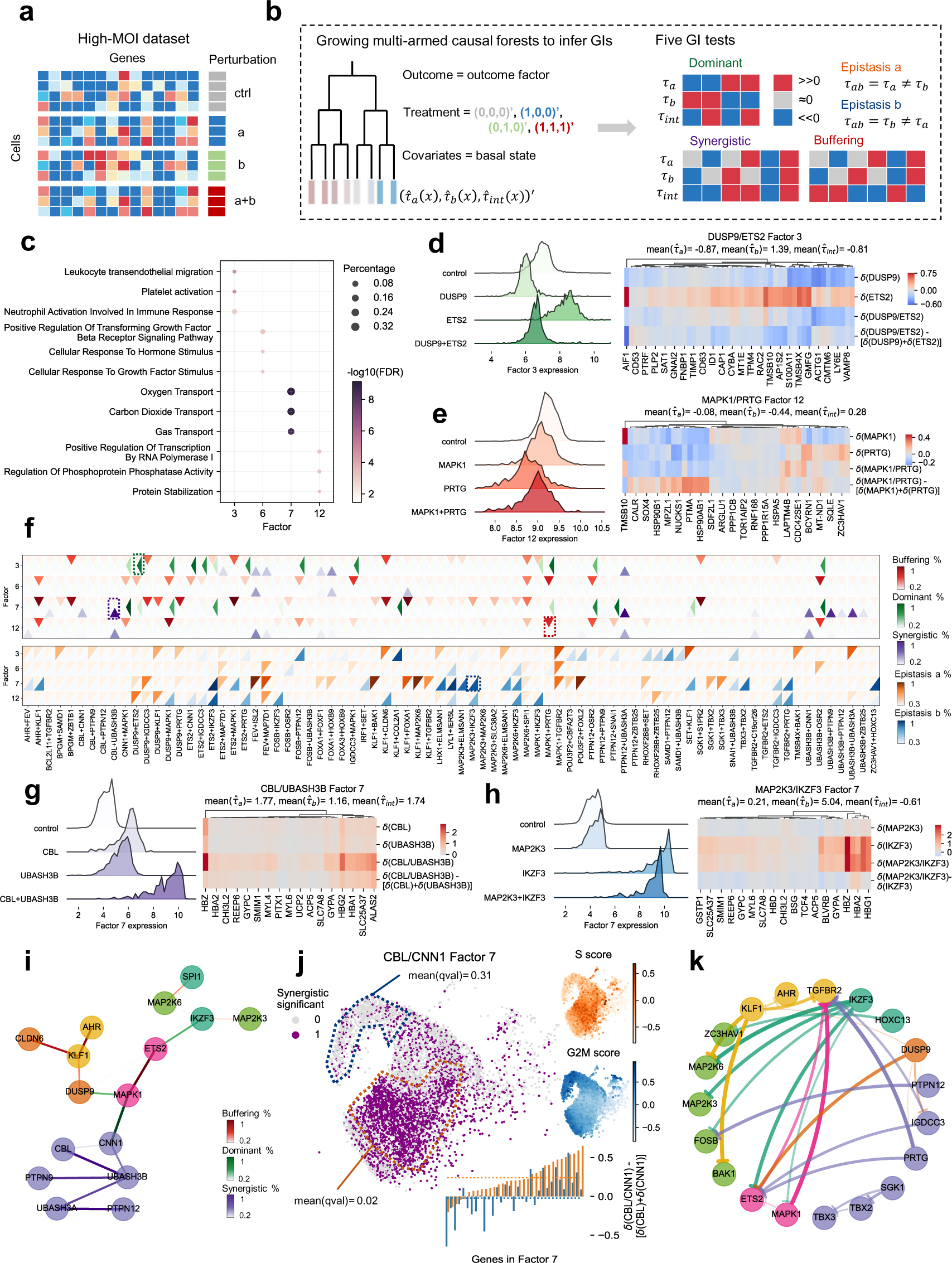
scCAPE infers genetic interactions in high-MOI datasets. (a) Illustration of a high-MOI (2-gene perturbation) single-cell CRISPR screening dataset. (b) The workflow of scCAPE in analyzing high-MOI (2-gene perturbation) single-cell CRISPR screening datasets. 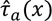: The perturbation effect estimation of perturbing gene *a* in cell *x*. 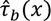: The perturbation effect estimation of perturbing gene *b* in cell *x*. 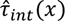: The estimation of interaction effects. ‘>>0 (<<0)’ means the effect is significantly positive (negative), and ‘≈0’ means the effect is not significant. (c) The gene set enrichment analysis of high-loading genes selected from factors 3, 6, 7 and 12. The size of the dot represents the percentage of selected genes in that Gene Ontology (GO) term^25^ or KEGG pathway^54^. The color of the dot shows the −log_10_(adjusted p-value). (d, e, g, h) The validation of four examples of GIs inferred by scCAPE. We compared the factor values of four conditions (left) and the expression of high-loading genes in that factor (right). *δ*(·) denotes average deviation in gene expression relative to control cells, thus *δ*(*a*/*b*) − [*δ*(*a*) + *δ*(*b*)] represents average interaction effects on these high-loading genes. (f) Results of genetic inference analysis (79 gene pairs) on factors 3, 6, 7, and 12. The color lightness indicates the percentage of cells that are significant (q-value<0.05) in that GI test. (i) The dominant, buffering, and synergistic interactions on factor 7 inferred by scCAPE. The color of the lines represents different types of GI. The width and lightness of the lines both indicate the percentage of cells that are significant (q-value<0.05) in the GI test. Genes with similar perturbation effects (see **Supplementary Figure 18**) are colored identically. (j) The heterogeneous synergistic effects in driving erythroid differentiation between CNN1 and CBL. We identified two subclusters (orange: synergy-significant enriched, blue: non-significant) using MELD^55^ and compared the average q-value. The interaction effects on high-loading genes in factor 7 (33 genes) averaged across these two subclusters are shown in the bar plot. The dotted lines represent the mean interaction effects across the 33 genes. (k) The epistasis interactions on the overall transcriptome inferred by scCAPE. Each arrow denotes an epistasis interaction originating in the gene that masks the effects of the other when the two genes are perturbed together. The width and lightness of the lines both represent the percentage of cells that are significant (combined q-value<0.05) in the epistasis test. Genes with similar perturbation effects (see **Supplementary Figure 18**) are colored identically.

For illustration, we applied scCAPE to identify genetic interactions from Perturb-seq data^20^ that studied 132 gene pairs in the K562 cell line. After preprocessing (**Methods**), we obtained the transcriptome profiles of 64,957 cells, consisting of 79 two-gene perturbations, their corresponding 55 one-gene perturbations, and untreated control. scCAPE selected 12 factors with distinct biological functions (**Figure 6c** and **Supplementary Figure 14**). We focus our discussion on factors 3, 6, 7, and 12. Factor 3 encompasses many megakaryocyte-related genes and exhibits enrichment for pathways of platelet activation and leukocyte transit. Factor 6 is associated with cellular response to pro-growth factors. Factor 7 has large loadings for erythroid markers such as hemoglobin genes. Factor 12 is enriched for GO terms related to the regulation of protein ubiquitination and phosphoprotein phosphatase activity.

The GI tests revealed diverse interaction effects between these gene pairs (**Figure 6f**), many of which have been documented in the literature. For instance, we found overexpression of DUSP9 and ETS2 yields opposing effects on factor 3, with DUSP9 exerting dominance in their combined perturbation (**Figure 6d**). MAPK1 and PRTG display buffering interactions on factor 12, where the effect of their combined perturbation is less than the sum of their individual perturbations (**Figure 6e**). CBL and UBASH3B demonstrate significant synergistic interactions in inducing erythroid markers (**Figure 6g**). IKZF3 and MAP2K3 exhibit an epistasis interaction on factor 7, with IKZF3 masking the effects of MAP2K3 when perturbing them together (**Figure 6h**).

A unique advantage of our approach is the evaluation of genetic interactions in a factor-specific manner, providing direct biological insights into these interactions. For instance, the gene UBASH3B, encoding ubiquitin-associated and SH3 domain-containing protein B, and PTPN9/12 (tyrosine phosphatases) are annotated in the GO term “peptidyl-tyrosine dephosphorylation”. scCAPE identified their interaction as synergistic, and only when factor 7 (erythroid differentiation) is considered as the outcome (**Figure 6f**). This highlights their shared mechanism in regulating tyrosine phosphorylation activity and promoting erythroid differentiation, consistent with previous studies^36–38^. To emphasize, scCAPE not only correctly identified the genetic interaction of UBASH3B and PTPN9/12, but also explicitly determined that the interaction affects erythroid differentiation. This specific information can guide subsequent validation experiments, leading to new insights. To explore the genetic interactions in driving erythroid differentiation, we utilized an undirected graph to visualize dominant, synergistic, and buffering interactions of selected genes regarding factor 7 (**Figure 6i**). For instance, we observed that overexpression of MAPK1, a mitogen-activated protein kinase involved in the development of megakaryocytes^39^, leads to decreased expression of erythroid genes. When perturbing MAPK1 and CNN1 (an erythroid differentiation driver) together, MAPK1 exerts dominance over CNN1, resulting in the combined perturbation antagonizing erythroid differentiation. In contrast, when perturbing MAPK1 and DUSP9 (a dual specificity phosphatase) together, DUSP9 dominates MAPK1 and induces erythroid markers. These findings align with the established biology of these pathways^40^.

Moreover, scCAPE reveals the heterogeneous landscape of genetic interactions. In other words, we can distinguish subpopulations of cells with and without interactions. For instance, our analysis identified synergy between CBL and CNN1 in promoting erythroid differentiation, as reported in the original study^20^. Furthermore, we revealed that this interaction was significant in 47% of cells, which were enriched in the S phase of the cell cycle (**Figure 6j and Supplementary Figure 15**). This additional finding is important as it pinpoints the cell cycle stage where the synergistic effect takes place. This discovery aligns with several prior studies suggesting that the transition of early erythroid progenitors to erythroid cells involves a swift transcriptional switch activating erythroid genes, and this switch synchronizes with and relies upon progression through the S phase^41^. Moreover, similar patterns were observed for other erythroid differentiation drivers (**Supplementary Figure 16**), such as PTPN12 and UBASH3B, UBASH3B and ZBTB25, etc. Therefore, our analysis implied that the S stage may set the context for the activation of transcription factors that are induced earlier in the progression of erythroid differentiation.

When the interest is to infer GIs at the transcriptome level instead of the specific factors, we provide an aggregated p-value that combines the p-values of different factors (**Methods**). **Figure 6k** illustrates the epistasis interactions on overall transcriptome concerning specific targets. For instance, simultaneous perturbation of IKZF3 (a B-cell transcription factor) and multiple genes in the MAP kinase signal transduction pathways (MAP2K3/6, MAPK1) results in a transcriptional profile akin to perturbing IKZF3 alone. When perturbing SGK1 (a serine/threonine-protein kinase) and genes in the T box transcription factor family (TBX2/3) together, the perturbation effects are masked by SGK1 (**Supplementary Figure 17**). These observations are consistent with the original study that investigated GIs at the whole transcriptome level^20^.

### Application of scCAPE in clustering perturbations

As a subsequent task, scCAPE can quantify the similarities between perturbations either in an overall manner or in functional factor-specific contexts. Additionally, benefiting from single-cell resolution estimations, these analyses can be conducted within any subpopulation of cells (**Figure 7a** and **Methods**).

**Figure 7.**
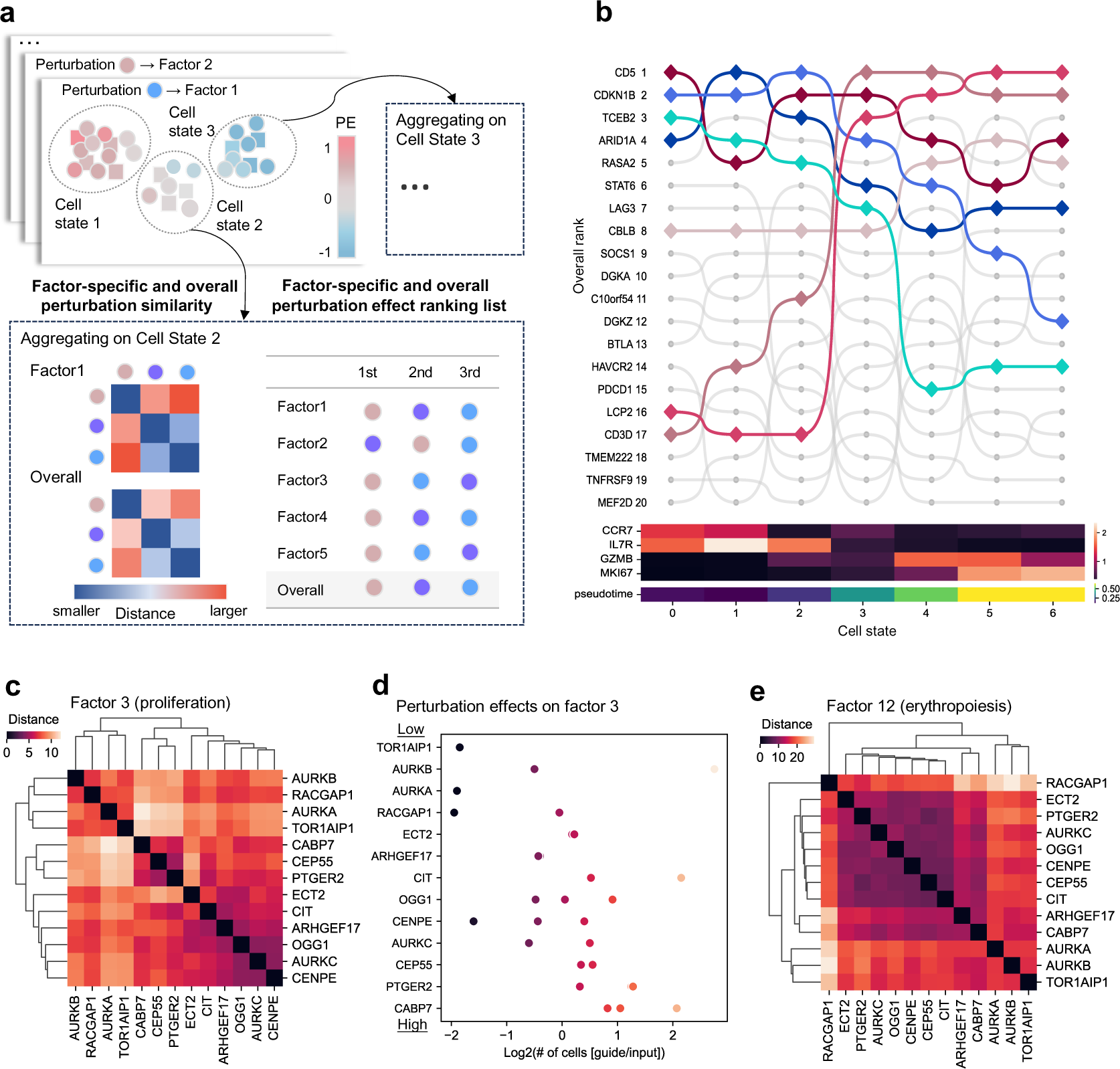
scCAPE quantifies the relationship between perturbations and prioritizes the perturbation effects. (a) Given the estimations of perturbation effects (PE) in each cell, scCAPE can further output the overall and factor-specific perturbation similarity and the ranking list at any cell subpopulation level. (b) The overall rankings of the perturbations in the CROP-seq T cell dataset along the TCR-stimulation trajectory. Initially, cells were clustered into 7 states, and a pseudotime trajectory was inferred to delineate the TCR-stimulation intensity (**Supplementary Figure 10**). Then, we calculated the overall perturbation ranking lists aggregated on 7 cell states, respectively. The heatmaps show the average expression of markers indicative of TCR-stimulated state (GZMB, MKI67) and unstimulated state (CCR7 and ILR7), and average pseudotime across these states. (c) The similarity between perturbations in the Perturb-seq K562 cell dataset on factor 3 (cell proliferation), aggregated on all cells. (d) The estimated perturbation effects on cell proliferation correlate with the observed fitness effects. Shown are the fold changes of gRNA abundance compared to their input abundance (x-axis) for the guides (dots) targeting each gene (y-axis). The targets are ranked by their perturbation effects on factor 3 (cell proliferation). (e) The similarity between perturbations in the Perturb-seq K562 cell dataset on factor 12 (erythropoiesis), aggregated on all cells.

The relationship between perturbations can vary across distinct cell states. For example, in the CROP-seq T cells dataset^11^, we investigated the clustering of perturbations on factor 7 (T cell proliferation). Within stimulated cells, we observed that CD3D and LCP2 exhibit similar positive effects on T cell proliferation, whereas genes such as RASA2, CBLB, TCEB2, SOCS1, CD5, and CDKN1B share similar negative effects on T cell proliferation (**Supplementary Figure 9a**). These findings align with the proliferation index of these genes^11^. However, in unstimulated cells, there is a slight alteration in the similarity pattern, with CBLB and SOCS1 no longer showing great similarity to the other four negative regulators, indicating that they might not act as general regulators of proliferation but instead modulate proliferation induced by TCR stimulation (**Supplementary Figure 9b**).

The clustering patterns of perturbations vary when looking at different outcome factors, thereby providing distinct biological insights. For instance, when examining factor-specific similarities among cell-cycle regulators within the Perturb-seq K562 cells dataset^3^, the distance matrix concerning factor 3 (cell proliferation) reveals two distinct clusters representing significant differences in proliferative outcomes (**Figure 7c**). In one cluster, the genes (CABP7, CEP55, and PTGER2) exert positive effects on factor 3, demonstrating a strong proliferative advantage, while the other cluster (AURKA/B, RACGAP1, and TOR1AIP1) exhibits a disadvantage. When conducting clustering analysis on factor 12 (erythropoiesis), the clustering pattern notably differs, with RACGAP1 being distinctly separated from TOR1AIP1 and AURKA/B (**Figure 7e**): the knockout of RACGAP1 decreases the expression of genes associated with erythropoiesis, consistent with prior studies indicating the pivotal role of RACGAP1 in erythrocyte biosynthesis^42^, whereas knockout of TOR1AIP1 and AURKA/B increases the expression of these erythropoiesis-related genes. Thus, by quantifying factor-specific perturbation effects, we elucidated the differing mechanisms of RACGAP1 and AURKA/B in regulating cell proliferation and erythroid differentiation. It is noteworthy that such results may be easily overlooked using transcriptome-level perturbation clustering methods (e.g., Euclidean distance between perturbation centroids in the PCA space^43^, E-distance^44^, etc.), as the number of genes in an informative factor may be very limited; for instance, factor 12 comprises only 21 high-loading genes, and the perturbation effect and subsequent clustering pattern may be overshadowed by modules or pathways comprising numerous genes.

When the interest is to quantify similarities at the transcriptome level, scCAPE provides the overall similarity matrix by aggregating results from all factors. We calculated the overall similarity among cell-cycle regulators within the Perturb-seq K562 cells dataset and observed that the aggregated clustering result of these perturbations predominantly aligns with their groupings by cellular morphology in HeLa cells^45^ (see **Supplementary Figure 19**).

### Application of scCAPE in prioritizing perturbation effects

Another application of scCAPE is to prioritize perturbations based on their effects (**Figure 7a** and **Methods**). The prioritizing can be conducted in an outcome factor-specific context. For example, we prioritized cell cycle regulators^3^ according to their effects on factor 3 (cell proliferation), and the ranking exhibited strong alignment with the observed fitness effects (**Figure 7d**).

Additionally, prioritization can be performed within subpopulations of cells to assess differences across distinct cell states. For instance, we examined the overall rankings of the perturbations in the CROP-seq T cells dataset^11^ along the TCR-stimulation trajectory (**Figure 7b** and **Supplementary Figure 10**). Notably, the relative importance of perturbations fluctuates in different cell states. The rankings of TCEB2 and CDKN1B decrease as the intensity of TCR stimulation escalates, whereas LCP2, CD3D, and CBLB exhibit an inverse trend, particularly for LCP2 and CD3D, their perturbation effects surge as cells transition from resting to stimulated states.

## Discussion

Single-cell CRISPR screening techniques have facilitated the efficient assessment of transcriptome-level impacts resulting from multiple genetic perturbations at single-cell resolution. While these methodologies present promising prospects, they also pose challenges in data analysis. Many current analytical approaches do not consider the confounding factors, potentially leading to biased perturbation effects estimation and ignoring the heterogeneity in perturbation effects. More complex methods that aim to predict the perturbation outcomes can suffer from issues related to interpretability^19,46–48^.

To tackle these challenges, we developed scCAPE to infer the heterogeneous perturbation effects at the single-cell level by framing the perturbation analysis as a problem of causal inference. scCAPE effectively separates the perturbation effects from the inherent cell-state variations and provides the nonparametric estimates and asymptotic results of perturbation effects on factors with distinct biological functions. Compared with other factor analysis methods^16,17^, the nonnegative and orthogonal constraints in scCAPE increase the factor interpretability and prevent the perturbation effects on different factors from canceling each other out. The cellular results in scCAPE empower us to examine the regulatory roles of the genetic perturbations in a more elaborated view and characterize subpopulations that respond differently to these perturbations. In subsequent tasks, scCAPE facilitates quantifying the relationship between perturbations and prioritizing them based on their overall regulatory impacts or their effects on specific functional factors.

Furthermore, scCAPE offers an approach for analyzing genetic interactions within high-MOI screening datasets, with three primary advantages over current methodologies^3,20,48^. Firstly, as a single-cell resolution method, scCAPE allows for the identification of heterogeneous interaction effects, which is not achievable with previous methods. Additionally, besides conducting transcriptome-level analyses like previous methods^20^, scCAPE examines interaction effects at the factor level, unveiling diverse types of genetic interactions within distinct pathways. Finally, it assesses genetic interactions by statistically testing the significance of interaction effects, eliminating the need for manual threshold settings on specific metrics^48^.

scCAPE is a versatile framework for single-cell data analysis and can be potentially extended to other settings. For example, it can be applied to drug-perturbation datasets to reveal heterogeneous treatment effects of the drugs across different cell states. There are several future directions for improvement. First, scCAPE can be enhanced to accommodate multi-omics perturbation analysis. Currently, besides scRNA-seq, single-cell epigenomic profiling (e.g., single-cell ATAC-seq)^49–51^ and the cell surface protein^52^ could serve as the screening readout, providing novel approaches to measure epigenome or proteome changes upon perturbation. It would be of interest to investigate the joint embedding and perturbation responses of data from multi-omics assays. Second, current inference results are conducted at the outcome factor level and are not straightforward to convert to the individual-gene level. Third, it would be desirable to design a formal statistical test to assess the existence of heterogeneity.

With the continuous advancement of single-cell CRISPR screening experiments, we foresee that scCAPE can serve as a valuable tool for gaining insights into the biological roles of genetic perturbations and examining their heterogeneous effects.

## Methods

### Data preprocessing

The preprocessing includes quality control and normalization. We removed low-quality cells that meet any of the following conditions: (1) the number of expressed genes is less than 200 or greater than 5000; (2) the percentage of mitochondrial genes is larger than 10%. Next, genes that are expressed in fewer than three cells are excluded. For normalization, the read counts of each gene are divided by the total counts of the corresponding cell and then multiplied by 10000, the resulting values are subject to logarithmic transformation. Finally, the top 6000 highly variable genes (HVG) are selected for subsequent analysis. In the high-MOI dataset^20^, if the cell counts of the two-gene perturbation or any of its two corresponding one-gene perturbations are less than 200, the cells with these perturbation labels were all removed. The above analysis was carried out with the Python package *scanpy*^56^.

### Autoencoder framework with adversarial training

In a CRISPR screening experiment, each cell is characterized by its normalized gene expression and perturbation label. For cell *i*, we denote its normalized gene expression vector as ***x***_*i*_, of *G* dimensions, and one-hot encoding of the perturbation label as ***s***_*i*_, such that *s*_*ij*_ = 1 if cell *i* has undergone the *j*-th type of perturbation and 0 otherwise. scCAPE takes as input data 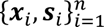 while *n* denotes the number of cells in the study.

An autoencoder with adversarial training was trained to map the original gene expression to a low-dimensional basal state (Equation 1).

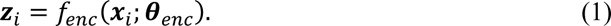

Next, a decoder was trained to generate gene expression levels from the basal state, by adding back the perturbation effects. In particular, we concatenated the perturbation label to the basal state (***z***_*i*_||***s***_*i*_), and passed forward to a neural network (Equation 2), and then mapped it back to the original gene expression space via factor analysis (Equation 3).

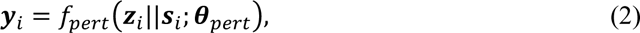

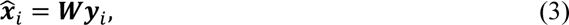

where ***W*** is a *G* × *K* non-negative orthogonal matrix, ***y***_*i*_ are *K*-dimensional outcome factors of gene expression.

The discriminator module predicts the perturbation label from the basal state (Equation 4), with binary cross-entropy loss (Equation 5).

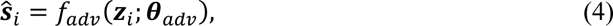

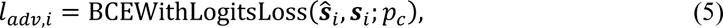

where *p*_*c*_ is the positive weight to adjust for the difference in cell numbers of perturbation labels.

In the training process, the discriminator is trained to predict the perturbation label ***s***_*i*_ given the basal state ***z***_*i*_, while the encoder is trained to make it impossible for the discriminator to predict ***s***_*i*_ given ***z***_*i*_ only. Specifically, for randomly selected minibatch data 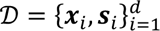 with *d* cells, the training minimizes the discriminator loss 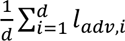 and the following loss (Equation 6) iteratively:

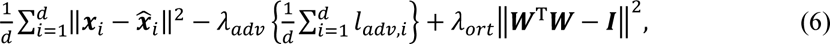

where *λ*_*adv*_ and *λ*_*ort*_ are hyperparameters to regularize the embedding space and the loading matrix. We set *λ*_*adv*_ = 1 and *λ*_*ort*_ = 0.5 if not stated otherwise. The detailed training algorithm is provided as **Algorithm 1 in Supplementary Note**.

The encoder *f*_*enc*_, perturbation network *f*_*pert*_ and discriminator *f*_*adv*_ are all fully connected neural network with ReLU activation, dropout and layer normalization. The dimensions of the layers and the values of tuning parameters in the networks are listed in **Supplementary Table 1**. We investigated the influence of varying hyper-parameters on the performance of scCAPE (**Supplementary Figure 20**), including the dimensions of the basal state, the coefficient of the adversarial loss (*λ*_*adv*_) and the coefficient of the orthogonal constraint (*λ*_*ort*_).

### Initialization of the loading matrix ***W***

We performed orthogonal non-negative matrix factorization^57,58^ (oNMF) for the original gene expression levels (Equation 7) to initialize ***W***. ***X***_*train*_ is the gene expression matrix of a subset of cells, randomly sampled from all cells in the dataset for computational efficiency.

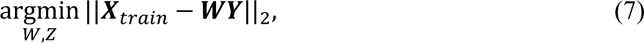

subject to ***W***^T^***W*** = *I*, *W*_*gk*_ ≥ 0 *Y*_*ki*_ ≥ 0, where *W*_*gk*_ denotes the loading of gene *g* on factor *k*, and *Y*_*ki*_ represents the value of *k*-th factor of cell *i*. Given the number of factors, the factorization can be solved through an iterative updating algorithm (**Algorithm 2 in Supplementary Note**).

The number of factors *K* should be pre-determined before running the oNMF algorithm. We choose *K* between 5∼15 to optimize the performance of the oNMF, which is evaluated using two criteria: (1) to minimize the reconstruction error (Equation 8); (2) to maximize the specificity score (*SS*_*k*_, Equation 9) for the factors, such that the loading of cells should differ as much as possible between perturbation groups.

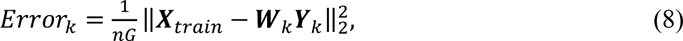

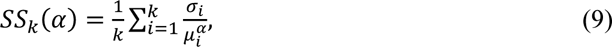

where *n* is the number of oNMF training samples (default 2,000), *G* is the number of HVGs in the dataset, *σ*_*i*_ and *μ*_*i*_ are the standard deviation and mean, respectively, of the *i*-th row of ***Y***_*k*_, *α* is a hyperparameter (default 1) controlling the power of the mean, when *α* = 1, the term in summation is the coefficient of variation.

We select the number of factors by minimizing *f*(*k*; *α j*), balancing the normalized version of the two scores (Equation 10):

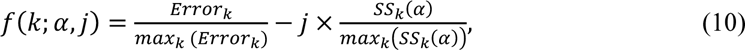

where *j* is the weight of the specificity score (default 0.3). Numerical implementation on the datasets **(Supplementary Figures 1, 6, 11 and 13)** shows that many hyperparameter settings can find a consistent set of the optimal number.

### Estimation of perturbation effects

We built causal forest models to estimate the cell specific perturbation effect under the potential outcome framework. Suppose the number of perturbation types is *S* and the number of factors is *K*, then for each combination of perturbation *s* (*s* ∈ {1 … *S*}) and outcome factor *k* (*k* ∈ {1 … *K*}), a causal forest is trained separately where the outcome is the factor (*y*_*ik*_) and the covariate is the basal state of a cell (***z***_*i*_). The treatment indicator is denoted as *t*_*i*_, where *t*_*i*_ = 1 if cell *i* has undergone perturbation *s* and 0 otherwise.

We consider the samples (***z***_*i*_, *y*_*ik*_, *t*_*i*_) ∈ ℝ^*D*^ × ℝ × {0, 1} with a random effect model (Equation 11)

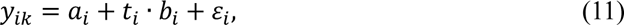

where *D* is the dimensions of the basal state, *ε*_*i*_ is the random noise. Define the intercept term *c*(***x***) = *E*[*a*_*i*_|***z***_*i*_ = ***x***] and the perturbation effect term *τ*(***x***) = *E*[*b*_*i*_|***z***_*i*_ = ***x***]. Under the unconfoundedness assumption {*a*_*i*_, *b*_*i*_, *ε*_*i*_} ⊥ *t*_*i*_|***z***_*i*_ (**Supplementary Note**), we have *τ*(***x***) is cell-specific perturbation effect that we aim to estimate. In addition, *τ*(***x***) can be identified by the local moment conditions through the following scoring function (Equation 12):

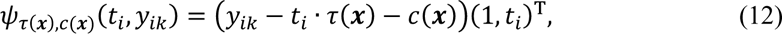

To estimate *τ*(***x***), we utilize the framework of generalized random forest^59^ which can efficiently infer the quantity identified via local moment conditions. Detailed discussion of the causal forest model and its estimation is provided in **Supplementary Note**.

Given the estimation and asymptotic variance at each cell, we can test whether there is a non-zero perturbation effect (**Algorithm 3 in Supplementary Note**). The q-values are then obtained from the FDR correction across all cells based on the function *statsmodels.stats.multitest. fdrcorrection* in the python package *statsmodels*^60^. The implementation of causal forests is based on the function *econml.grf.CausalForest* in the python package *econml*^61^.

### Annotating the functions of factors

To interpret the latent factors, we perform gene set enrichment analysis. For each factor, genes with weights greater than 2 times the standard deviation (*>* 2*σ*) are selected to perform the functional enrichment annotation to GO ‘biological process’ category^25^ and/or KEGG pathways^54^ using the *enrichr* module in a python package named *GSEApy*^62^.

### Genetic interaction analysis

We build an interaction causal forest (ICF) to infer genetic interactions (GIs) of perturbations. ICF is also implemented in a causal forest framework, while the treatment levels correspond to the combinations of perturbation status. Consider two perturbations *a* and *b*, a treatment vector (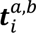) denotes the perturbation status of cell *i*, where (0,0,0)^T^, (1,0,0)^T^, (0,1,0)^T^ and (1,1,1)^T^correspond to control cells, cells with perturbation *a* only, cells with perturbation *b* only, and cells with both perturbations *a* and *b*. Based on this treatment coding, the quantity of interest ***τ***(***x***) = (*τ*_*a*_(***x***), *τ*_*b*_(***x***) *τ*_*int*_(***x***)) becomes a triplet which characterizes the perturbation effect of perturbing gene *a*, the perturbation effect of perturbing gene *b* and the interaction term, respectively.

Four types of GIs are considered: buffering, dominant, synergistic and epistasis^3,20^. We first test the main effect of each perturbation:

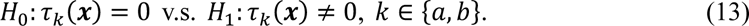

For cells with opposing single-gene perturbation effects, the dominant interaction effect is tested:

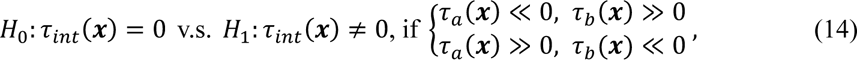

where *τ*_*k*_(***x***) ≪ 0 (≫ 0) means *τ*_*k*_(***x***) is significantly less (more) than zero, *k* ∈ {*a b*}.

The synergistic interaction tests are different according to the sign of the perturbation effects:

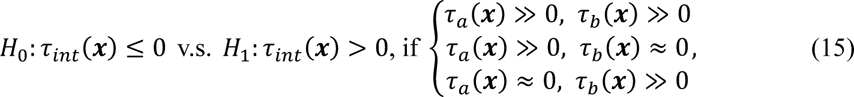

and

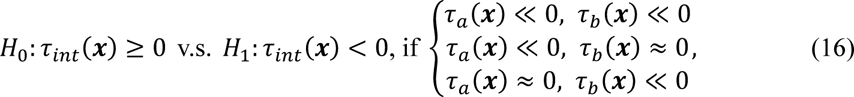

where *τ*_*k*_(***x***) ≈ 0 means *τ*_*k*_(***x***) is not significant (the q-value of the main effect test is larger than 0.05), *k* ∈ {*a,b*}.

The buffering interaction tests are

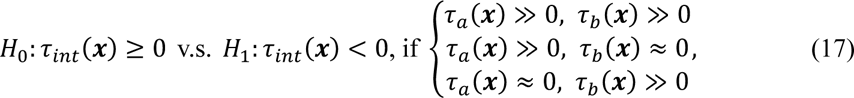

and

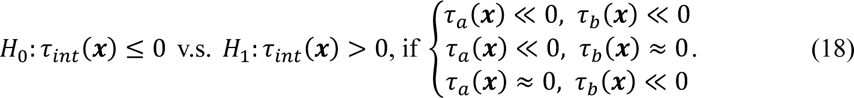

The epistasis interaction tests are conducted to check if one perturbation masks the effect of the other one in the combined perturbation:

1. Epistasis *a* test: *H*_0_: *τ*_*b*_(***x***) + *τ*_*int*_ (***x***) = 0 v.s. *H*_1_: *τ*_*b*_(***x***) + *τ*_*int*_(***x***) 0,
2. Epistasis *b* test: *H*_0_: *τ*_*a*_(***x***) + *τ*_*int*_(***x***) = 0 v.s. *H*_1_: *τ*_*a*_(***x***) + *τ*_*int*_(***x***) 0,

then if the effects of the combined perturbation are similar to that of gene-*a* perturbation while different from gene-*b* perturbation, the null hypothesis in (1) would be accepted and the null hypothesis in (2) would be rejected.

To give a genetic interaction inference on the overall transcriptome, we combine p-values of the GI tests on different factors using the aggregated Cauchy association test (ACAT)^63^, since ACAT accounts for arbitrary correlation structures between tests.

### Factor-specific and overall perturbation similarity

For each factor, we can quantify the similarity of perturbation effects in a subpopulation of cells (*C*). Denote the similarity matrix of factor *k* as ***D***_*k*_(*C*), whose *i j*-th entry is

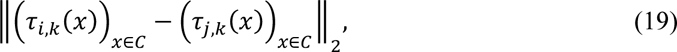

where *τ*_*i*_,_*k*_(*x*) is the perturbation effect of perturbation *i* on factor *k* in cell *x*.

An overall similarity matrix ***D***(*C*) is calculated by the weighted sum of factor specific similarity matrices:

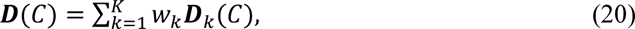

where *w*_*k*_ is the weight of factor *k*, which is proportional to the number of the genes with large loadings (*>* 2*σ*), and 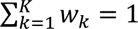.

### Factor-specific and overall perturbation effect ranking list

We prioritize the perturbations in a factor-specific way to explore which perturbations have large effects on these factors. In a subpopulation of cells (*C*), the perturbation effect ranking list w.r.t. factor *k* is *R*_*k*_(*C*), which is ranked by their average perturbation effects on *C*:

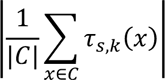

The overall ranking list *R*(*C*) is obtained by aggregating the ranking lists of different factors *R*_*k*_(*C*) with the weights of factors using R package *RankAggreg*^64^.

### Estimation of perturbation effects at gene level

The estimation of perturbation effects at the original gene expression level can be derived by linear transformation:

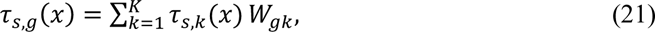

where *τ*_*s*_,_*g*_(*x*) denotes the perturbation effect of *s* on gene *g* in cell *x*.

### Simulation

We simulated a Perturb-Seq dataset based on the PBMC dataset^22^. Initially, we removed stimulated cells, and manually labeled the control cells to make the cell types have clear separation (**Supplementary Figure 3**). Then 50% of the control cells were randomly selected as stimulated cells, by introducing perturbation effects on factor level. Specifically, we added perturbation effect to factor 14 at varying levels, and then transformed the factor level expression back to the original gene expression level (Equation 22).

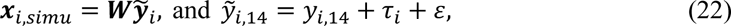

where *τ*_*i*_ denotes the varying perturbation effect on different cell types, *τ*_*i*_ = 0,0.1,0.2,0.3,0.4,0.5 or 0.6 if the cell type of cell *i* is CD4-T, CD8-T, NK, B, DC, F-Mono or CD14-Mono, and *ε* is a Gaussian noise with *σ* = 0.01. These effect sizes chosen at levels comparable to existing studies^17^. We generated 100 datasets and examined estimations and inferences of stimulation effects on factor 14 averaged across the seven cell types.

### Implementation of alternative methods

To demonstrate the disentangling performance of our method, we applied the following available methods for a comparison: (1) Mixscape^24^: We calculated the embedding for each stimulated cell by averaging the expression of its nearest-*k* control neighbors. *k* is set to be 20 as the paper stated. (2) CINEMA-OT^23^: a method achieves disentanglement via independent component analysis and optimal transport (OT), using the function *cinemaot.cinemaot_unweighted* in the Python package *cinemaot*. (3) OT: classical optimal transport, using the function *SinkhornKnopp* in the python package *cinemaot*.

For comparing the estimations of average perturbation effects, we applied the following perturbation analysis method when analyzing the Perturb-seq K562 cells dataset: (1) MIMOSCA^3^: We fit a linear regression model between the gene expression matrix and the perturbation labels using the function *sklearn.linear_model.ElasticNet(precompute=True, l1_ratio=0.5, alpha=0.0005, max_iter=10000)* as the paper stated. (2) MIMOSCA+ cell state: We fit a linear regression model between the gene expression and the perturbation labels with pre-determined 9 cell states using *sklearn.linear_model.ElasticNet(precompute=True,l1_ratio=0.5,alpha=0.0005,max_iter=10000)* as the paper stated.

### Evaluation metrics

Five evaluation metrics are employed to evaluate the mixing performance of disentangling methods, including ASW^65^, PCR^66^, LISI^67^, kBET^66^, and XGBoost^68^. For ASW, PCR, LISI and kBET, the metrics are all adjusted to maximum 1 with the approach stated in ref.69, and the larger score means better mixing performance. For XGBoost, we split the data into training and test sets and fit a XGBoost model to predict the perturbation labels based on the embedding of the training samples. We calculate the prediction accuracy in the test set, and use the percentage of misclassification as an evaluation metric since lower accuracy indicates better mixing performance. The implementation of XGBoost is based on the R package *xgboost*.

## Data availability

All of the datasets analyzed in this manuscript are publicly available. The PBMC data were accessed from scGen’s tutorial^47^, with the original data from ref.22 under GSE96583. The T cells CROP-seq data were taken from the original publication^11^ under GSE119450. The K562 Perturb-seq data^3^ were accessed from the Single Cell Portal https://singlecell.broadinstitute.org/single_cell/study/SCP31/perturb-seq-k562. The high-MOI data were accessed from ref.20 under GSE133344.

## Code availability

Our method is implemented as an open-source Python package available at https://github.com/zcfu21/scCAPE.

## Notes

### Competing Interest Statement

The authors have declared no competing interest.

