## Supplementary Material for "Inference of heterogeneous effects in single-cell genetic perturbation screens"

### 1 Supplementary Notes

#### 1.1 Treatment effects estimation, CATE

We formulate the perturbation analysis as the problem of conditional average treatment effect estimation (CATE) in causal inference[1,2]. Consider the perturbation as 'treatment', denoted by $T_i \in \{0,1\}$, where $T_i = 1$ if cell $i$ is perturbed, and $T_i = 0$ otherwise. The transcriptomic responses to the perturbation (outcome factors) is the 'outcome' variable, denoted by $Y_i$. The covariates, such as inherent cell status (basal state), are denoted as $X_i \in \mathbb{R}^d$. Following the potential outcome framework[1,2], there exists potential outcomes $Y_i(1)$ and $Y_i(0)$ corresponding respectively to the response the $i$-th cell would have experienced with and without the perturbation. Our interest is to estimate the CATE, which is averaged on a subpopulation or a single cell:

$$\tau(x) = \mathbb{E}[Y_i(1) - Y_i(0) | X_i = x]. \tag{1}$$

Under the assumption of unconfoundedness[3] CATE can be identified as a statistical estimand:

$$\tau(x) = \mathbb{E}[Y | T = 1, X = x] - \mathbb{E}[Y | T = 0, X = x]. \tag{2}$$

#### 1.2 Causal forests

There have been extensive methods to estimate the conditional average treatment effects under the assumption of unconfoundedness[3], including classical methods like (generalized) linear models, nearest-neighbor matching, kernel methods, series estimation (see e.g. refs[4-6]), causal forests[7-9] and neural network methods (see a review in ref[10]). In scCAPE, we adopt the framework of generalized random forest[8] (GRF), since the method is adaptive, non-parametric, and provides estimations with asymptotic properties.

The rationale behind causal forests is that samples with similar covariates can be treated as though they had come from a randomized experiment. Thus, the covariate space can be adaptively partitioned into small 'leaves', within which the outcomes of control and treatment samples can be compared to estimate treatment effects. Below is an overview of the causal forest procedures. For a single causal tree, a subset of samples is randomly selected, and the covariate space is adaptively partitioned by creating splits that maximize treatment effect heterogeneity. Splitting continues until the tree reaches a pre-determined depth or each leaf contains no more than a specified minimum number of control or treated samples ($n_{min}$). Within each leaf, the treatment effects are estimated by comparing outcomes between treated and control samples. Given the procedure for generating a single causal tree, a causal forest creates numerous such trees and then aggregates their estimations. In addition, the asymptotic variances of the estimates can be analyzed using theorems of U-statistics[8].

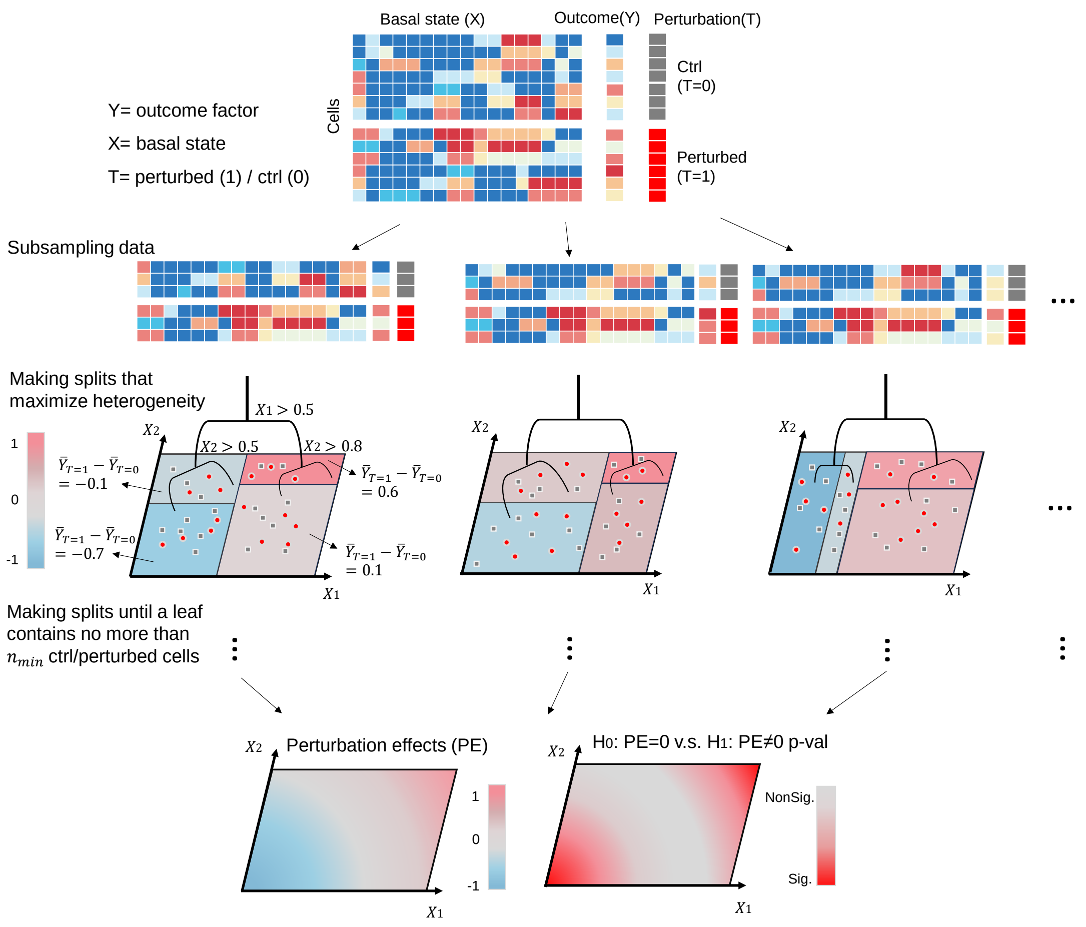

Here we consider the samples of cells $(X_i, Y_i, T_i) \in \mathbb{R}^d \times \mathbb{R} \times \{0,1\}$ with a random effect model

$$Y_i = a_i + T_i \cdot b_i + \varepsilon_i, \tag{3}$$

where $a_i$ and $b_i$ are random effects, and $\varepsilon_i$ is a random noise with $\mathbb{E}[\varepsilon_i|X_i = x] = 0$. Define the intercept term (nuisance parameter) $c(x) = \mathbb{E}[a_i|X_i = x]$ and the perturbation effect term $\tau(x) = \mathbb{E}[b_i|X_i = x]$. In addition, we assume that $\{a_i, b_i, \varepsilon_i\} \perp T_i|X_i$, which is equivalent to the unconfoundedness[3]. Under the assumption, we have $\mathbb{E}[Y_i|T_i, X_i = x] = c(x) + T_i \cdot \tau(x)$, which indicates $\tau(x)$ is exactly the CATE we aim to estimate*. To estimate $\tau(x)$, we can use the following scoring function:

$$\psi_{\tau(x),c(x)}(T_i, Y_i) = (Y_i - T_i \cdot \tau(x) - c(x))(1, T_i)^{\mathrm{T}}, \tag{4}$$

$\tau(x)$ can be identified by the local moment conditions $\mathbb{E}\left[\psi_{\tau(x),c(x)}(T_i, Y_i)|X_i = x\right] = 0$ through the above scoring function and be written explicitly as**

$$\tau(x) = \{\mathrm{Var}[T_i|X_i = x]\}^{-1}\mathrm{Cov}[T_i, Y_i|X_i = x]. \quad (5)$$

GRF estimates $\tau(x)$ by growing causal forests. In the process of growing trees, GRF adopts a subsample splitting technique called honesty to eliminate bias[9] and uses a gradient-based approximation splitting rule to maximize heterogeneity of $\tau$ between the two branches. Suppose GRF grows a set of $B$ trees indexed by $b = 1, \dots, B$ and, for each tree, defines $L_b(x)$ as the set of training samples falling in the same "leaf" as $x$. The forests weights $\alpha_i(x)$ can be calculated from the frequency with which the $i$-th training sample falls into the same leaf as $x$:

$$\alpha_{bi}(x) = \frac{1(X_i \in L_b(x))}{|L_b(x)|}, \quad \alpha_i(x) = \frac{1}{B}\sum_{b=1}^{B}\alpha_{bi}(x). \quad (6)$$

Given forests weights $\alpha_i(x)$ defined in Eq.(6), the estimator is

$$\hat{\tau}(x) = \left[\sum_{i=1}^{n}\alpha_i(x)(T_i - \bar{T}_\alpha)^2\right]^{-1}\sum_{i=1}^{n}\alpha_i(x)(T_i - \bar{T}_\alpha)(Y_i - \bar{Y}_\alpha), \quad (7)$$

where $\bar{T}_\alpha = \sum\alpha_i(x)T_i$ and $\bar{Y}_\alpha = \sum\alpha_i(x)Y_i$.

The estimation of asymptotic variances of $\hat{\tau}(x)$ can be derived by estimating the derivatives of the scoring function and the variances of weighted scoring function, which can be seen in ref.8.

For interaction causal forests (ICF) defined in the paper, with the same procedures in growing trees and calculating forest-based weights $\alpha_i(x)$, $\boldsymbol{\tau}(x)$ can be estimated by

$$\hat{\boldsymbol{\tau}}(x) = \left[\sum_{i=1}^{n}\alpha_i(x)(\boldsymbol{T_i} - \bar{\boldsymbol{T}}_{\boldsymbol{\alpha}})^{\otimes 2}\right]^{-1}\sum_{i=1}^{n}\alpha_i(x)(\boldsymbol{T_i} - \bar{\boldsymbol{T}}_{\boldsymbol{\alpha}})(Y_i - \bar{Y}_\alpha), \quad (8)$$

where $\bar{\boldsymbol{T}}_{\boldsymbol{\alpha}} = \sum\alpha_i(x)\boldsymbol{T_i}$ and $\bar{Y}_\alpha = \sum\alpha_i(x)Y_i$, and $w^{\otimes 2} = ww^{\mathrm{T}}$. The asymptotic property still holds as shown in ref.8.

[Derivations of the claims:]

*$\mathbb{E}[Y_i|T_i, X_i = x] = \mathbb{E}[a_i + T_i \cdot b_i + \varepsilon_i|T_i, X_i = x] = \mathbb{E}[a_i|T_i, X_i = x] + \mathbb{E}[T_i \cdot b_i|T_i, X_i = x] + \mathbb{E}[\varepsilon_i|T_i, X_i = x] = \mathbb{E}[a_i|X_i = x] + T_i \cdot \mathbb{E}[b_i|X_i = x] + \mathbb{E}[\varepsilon_i|X_i = x]$(unconfoundedness assumption) $= c(x) + T_i \cdot \tau(x)$, and thus

$$\tau(x) = \mathbb{E}[Y|T = 1, X = x] - \mathbb{E}[Y|T = 0, X = x],$$

which means $\tau(x)$ is exactly the CATE we aim to estimate.

** $\mathrm{Cov}[T_i, Y_i|X_i = x] = \mathrm{Cov}[T_i, T_i b_i|X_i = x] = \mathbb{E}\left[T_i^2 b_i|X_i = x\right] - \mathbb{E}[T_i|X_i = x]\mathbb{E}[T_i b_i|X_i = x] = \mathbb{E}\left[T_i^2|X_i = x\right] \cdot \tau(x) - (\mathbb{E}[T_i|X_i = x])^2 \cdot \tau(x)$ (unconfoundedness assumption) $= \mathrm{Var}[T_i|X_i = x] \cdot \tau(x)$, which gives Eq.(5). To see $\tau(x)$ is identified by the scoring function defined in Eq.(4), we can solve $\mathbb{E}\left[\psi_{\tau(x),c(x)}(T_i, Y_i)|X_i = x\right] = (0,0)^{\mathrm{T}}$:

$$\mathbb{E}\left[\psi_{\tau(x),c(x)}(T_i, Y_i)|X_i = x\right] = \begin{bmatrix} \mathbb{E}[Y_i|X_i = x] - \tau(x)\mathbb{E}[T_i|X_i = x] - c(x) \\ \mathbb{E}[Y_i T_i|X_i = x] - \tau(x)\mathbb{E}\left[T_i^2|X_i = x\right] - c(x)\mathbb{E}[T_i|X_i = x] \end{bmatrix} = \begin{bmatrix} 0 \\ 0 \end{bmatrix},$$

we have $\tau(x) = \{\mathrm{Var}[T_i|X_i = x]\}^{-1}\mathrm{Cov}[T_i, Y_i|X_i = x]$, which yields the conclusion.

#### 1.3 Algorithm of adversarial training

**Algorithm 1:** Mini-Batch training of scCAPE

**Input:** Data matrix $\boldsymbol{X}$ with HVGs, one-hot encoding of perturbation label $s$ of each cell, batch size $B$, training epochs $n_{\text{epoch}}$, discriminator training steps $n_{adv}$, the weight of the discriminator loss $\lambda_{adv}$, the weight of the orthogonal loss $\lambda_{ort}$.

**Output:** The perturbation-invariant latent representation $\boldsymbol{z}$ and the factor-level expression $\boldsymbol{y}$ of each cell. The non-negative orthogonal loading matrix $\boldsymbol{W}$.

Initialize the weights of networks in scCAPE. Initialize the non-negative orthogonal loading matrix $\boldsymbol{W}$ (see Algorithm 2). Set iteration times $t = 0$.
**for** $i = 1$ **to** $n_{\text{epoch}}$ **do**
    Discriminator loss weight ($\lambda_{adv}$) scheduling
    **for** $b = 1$ **to** $\lfloor n/B \rfloor$ **do**
        $t \leftarrow t + 1$, sample $B$ cells from the training dataset
        **if** $t \,\%\, n_{adv} == 0$ **then**
            Minimize the discriminator loss $\boldsymbol{\theta}_{adv} \leftarrow \underset{\boldsymbol{\theta}_{adv}}{\operatorname{argmin}}\, l_{adv}$
        **else**
            Minimize the loss
            $\boldsymbol{\theta}_{enc}, \boldsymbol{\theta}_{pert}, \boldsymbol{W} \leftarrow \underset{\boldsymbol{\theta}_{enc}, \boldsymbol{\theta}_{pert}, \boldsymbol{W}}{\operatorname{argmin}} \left(l_r - \lambda_{adv} l_{adv} + \lambda_{ort} ||\boldsymbol{W}^{\mathrm{T}}\boldsymbol{W} - \boldsymbol{I}||^2\right)$
        **end if**
    **end**
**end**
**return** $\boldsymbol{z}$, $\boldsymbol{y}$ and $\boldsymbol{W}$

$l_r$ is the MSE reconstruction loss. Following the suggestion of ref [11], we initially set $\lambda_{adv}$ to 0 and the model is trained like a normal autoencoder. Then, $\lambda_{adv}$ is linearly increased to the set value over the first fourth of the maximum training epochs to slowly encourage the model to produce invariant representations.

#### 1.4 Algorithm of initialization of the loading matrix

The following algorithm is based on refs.12,13.

**Algorithm 2:** Orthogonal nonnegative matrix factorization

**Input:** Data matrix $\boldsymbol{X}$, factor number $K$, iteration times $m$ and convergence criterion $\epsilon$.

**Output:** The non-negative orthogonal loading matrix $\boldsymbol{W}$ and the latent factor matrix $\boldsymbol{Y}$.

Initialization of $\boldsymbol{W}$ and $\boldsymbol{Y}$. Perform K-means clustering of columns of $\boldsymbol{X}$. From this we obtain the cluster centroids $\boldsymbol{C} = (\boldsymbol{c}_1, \cdots, \boldsymbol{c}_K)$ and set $\boldsymbol{W} = \boldsymbol{C}$. From the cluster memberships, we obtain the factors $\boldsymbol{Y}$. Set $\boldsymbol{Y} \leftarrow \boldsymbol{Y}$+0.2.

**for** $t = 1$ **to** $m$ **do**

$$Y_{ki} \leftarrow Y_{ki} \frac{(W^{\mathrm{T}}X)_{ki}}{(W^{\mathrm{T}}WY)_{ki}},$$

$$W_{gk} \leftarrow W_{gk} \frac{(XY^{\mathrm{T}})_{gk}}{(WW^{\mathrm{T}}XY^{\mathrm{T}})_{gk}}$$

**if** converge **then**

**break**

**end if**

**end**

**return** $\boldsymbol{W}$ and $\boldsymbol{Y}$

#### 1.5 Algorithm of causal forests

The following algorithm is based on ref.8.

---

**Algorithm 3:** Inference of CATE using causal forests

---

**Input:** For a given perturbation and a factor $k$, collecting the set of examples $\mathcal{S} = \{\boldsymbol{z}_i,\ t_i,\ y_{ik}\}$, where $\boldsymbol{z}_i$ is the basal state, the treatment $t_i = 0/1$ if the cell is control/perturbed and $y_{ik}$ is the factor level expression (the corresponding element of $\boldsymbol{y}_i$). Tunning parameters like the number of trees $B$, subsampling rate $s$, etc.

**Output:** The CATE estimates and the corresponding q-value to the test $H_0\colon \mathrm{CATE} = 0$ of each cell $x$.

1. weight vector $\alpha \leftarrow$ ZEROS($|\mathcal{S}|$)
2. **for** $b = 1$ **to** $B$ **do**
3.     set of examples $\mathcal{I} \leftarrow$ SUBSAMPLE($\mathcal{S}, s$)
4.     set of examples $\mathcal{J}_1, \mathcal{J}_2 \leftarrow$ SPLITSAMPLE($\mathcal{I}$) [Honesty]
5.     tree $\mathcal{T} \leftarrow$ GRADIENTTREE($\mathcal{J}_1$)
6.     neighborhood $\mathcal{N} \leftarrow$ NEIGHBORS($x, \mathcal{T},\ \mathcal{J}_2$)
7.     **for** all example $e \in \mathcal{N}$ **do**
8.         $\alpha[e] \mathrel{+}= 1/|\mathcal{N}|$
9. calculate the CATE estimate $\hat{\tau}(x)$ from Eq.(7) with weights $\alpha/B$
10. $\hat{\sigma}^2(x) \leftarrow$ CALCULATEVAR
11. obtain p-value $P(|z| > |\hat{\tau}(x)/\hat{\sigma}(x)|)$ of each cell and do FDR correction across all cells
12. **return** $\hat{\tau}(x)$ and q-value of each cell

---

The function ZEROS creates a vector of zeros of length $|\mathcal{S}|$; SUBSAMPLE selects a subsample of size $s$ from $\mathcal{S}$ without replacement; SPLITSAMPLE randomly separates a set into two evenly sized, nonoverlapping halves; GRADIENTTREE grows a tree by running CART splits iteratively based on the pseudo-outcomes approximated by the gradient to maximize heterogeneity; NEIGHBORS returns those elements of $\mathcal{J}_2$ that fall into the same leaf as $x$ in the tree $\mathcal{T}$; CALCULATEVAR outputs the asymptotic variance based on a bootstrap-of-little-bags approach with an objective Bayesian debiasing correction.

---

#### 2 Supplementary Tables

**Supplementary Table 1. Hyperparameters for scCAPE**

| | Hyperparameter | Default value |
|---|---|---|
| General | Embedding dimension (basal state) | 20 |
| | Batch size | 256 |
| | Learning rate decay schedule, in epochs | 45 |
| | Weight of adversarial loss | 1, with increasing scheduling |
| | Weight of orthogonal constraint | 0.5 |
| Encoder | Layer dimension | [512, 128, 64] |
| | Learning rate | 3e-4 |
| | Weight decay | 4e-7 |
| Perturbation NN | Layer dimension | [20, 20, 20] |
| | Learning rate | 3e-4 |
| | Weight decay | 4e-7 |
| Discriminator | Layer dimension | [64, 64] |
| | Learning rate | 3e-4 |
| | Weight decay | 4e-7 |
| | Training steps | 3 |

#### 3 Supplementary Figures

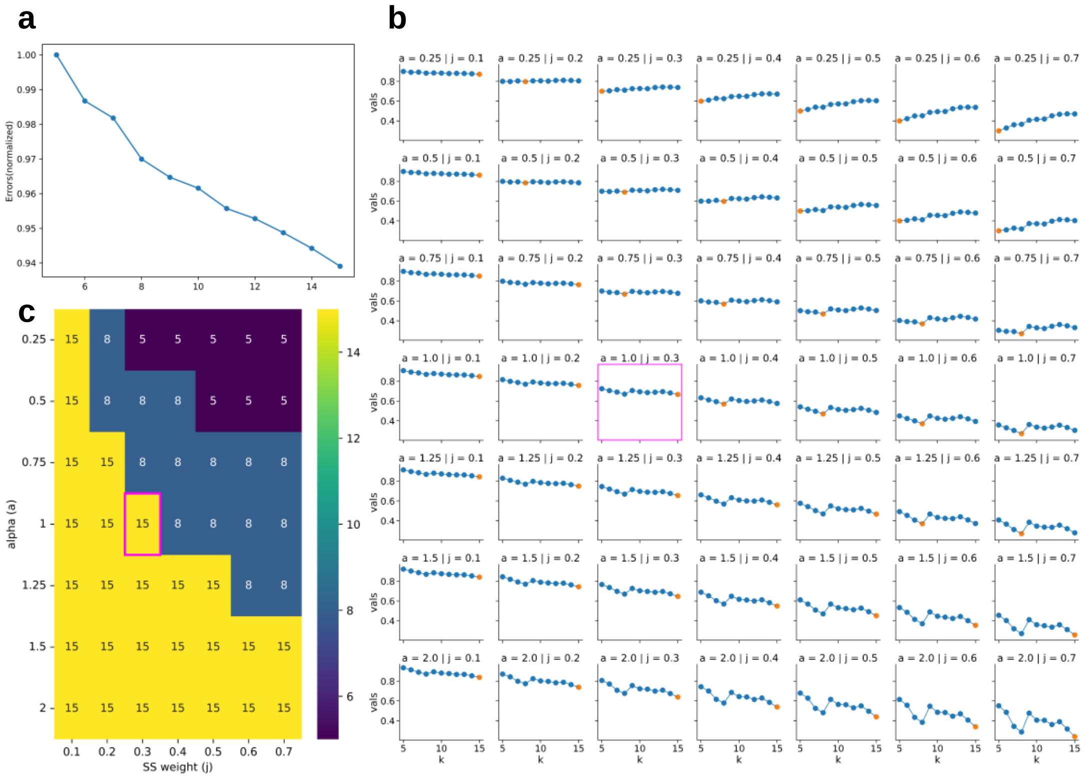

**Supplementary Figure 1. Selection of the number of factors in the PBMC dataset.**

(a) Normalized reconstruction errors across different factor numbers.

(b) The value of $f(k;\alpha,j)$ for a varying range of hyperparameter $\alpha$ (rows) and the weight of specificity score $j$ (columns). The highlighted points represent the minimum values. The bounding box shows the curve under the default setting ($\alpha = 1, j = 0.3$).

(c) The optimal number of factors under varying $\alpha$ and $j$. The bounding box shows 15 is the optimal number under the default setting ($\alpha = 1, j = 0.3$).

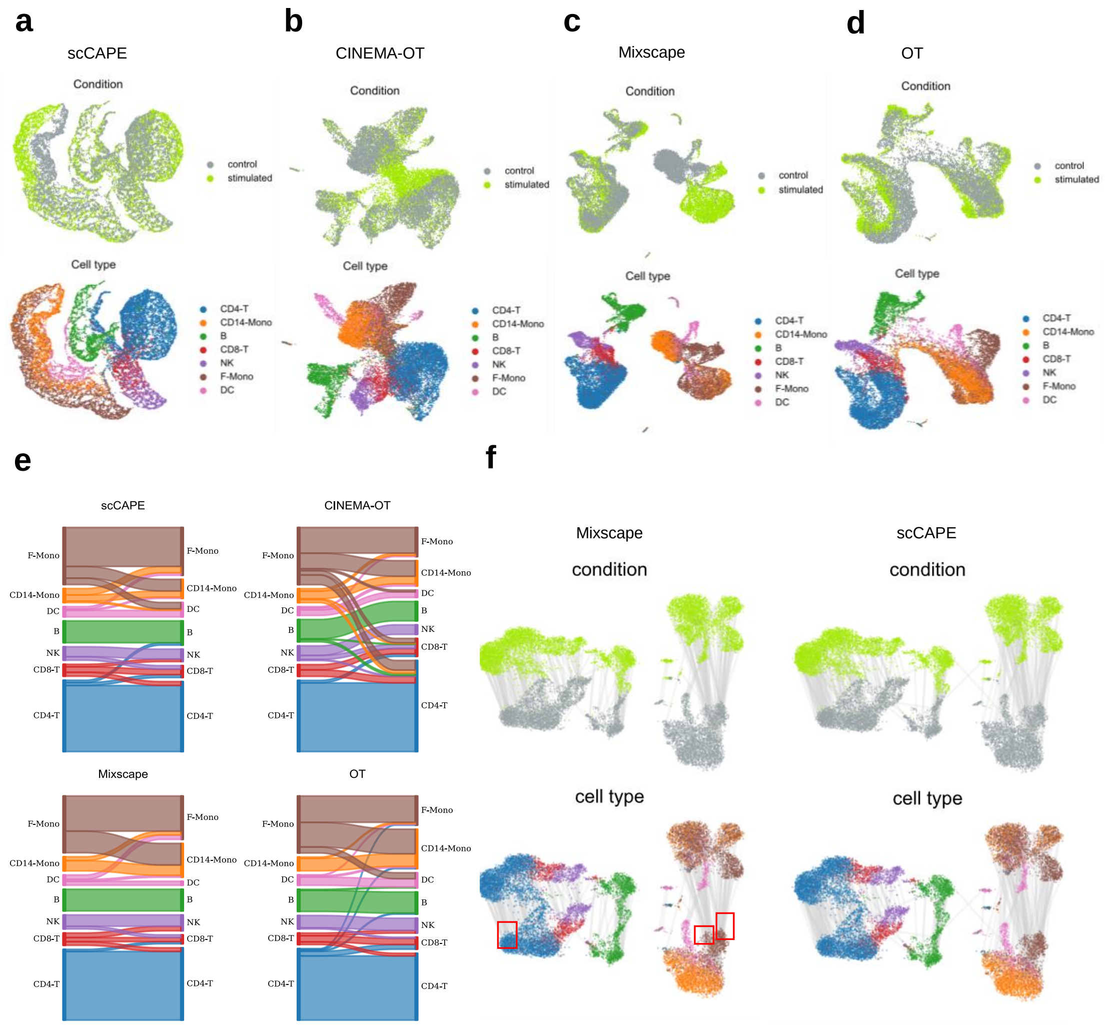

**Supplementary Figure 2. Visualizations of the disentangling results in the PBMC dataset.** (a-d) UMAP visualizations of the basal state, colored by conditions (top) and cell types (bottom). (e) The cell type of the nearest control neighbor for each stimulated cell in the embedding space of scCAPE, CINEMA-OT, OT and Mixscape. (f) UMAP visualizations of the original data, colored by conditions (top) and cell types (bottom). Each stimulated cell and its nearest control neighbor in the latent embedding (basal state in Mixscape and scCAPE) are connected by a line. In Mixscape, many stimulated cells are matched with the same group of control cells on the boundary of clusters (red boxes). As a global method that minimizes the loss function over all cells, scCAPE alleviates this “boundary effects”.

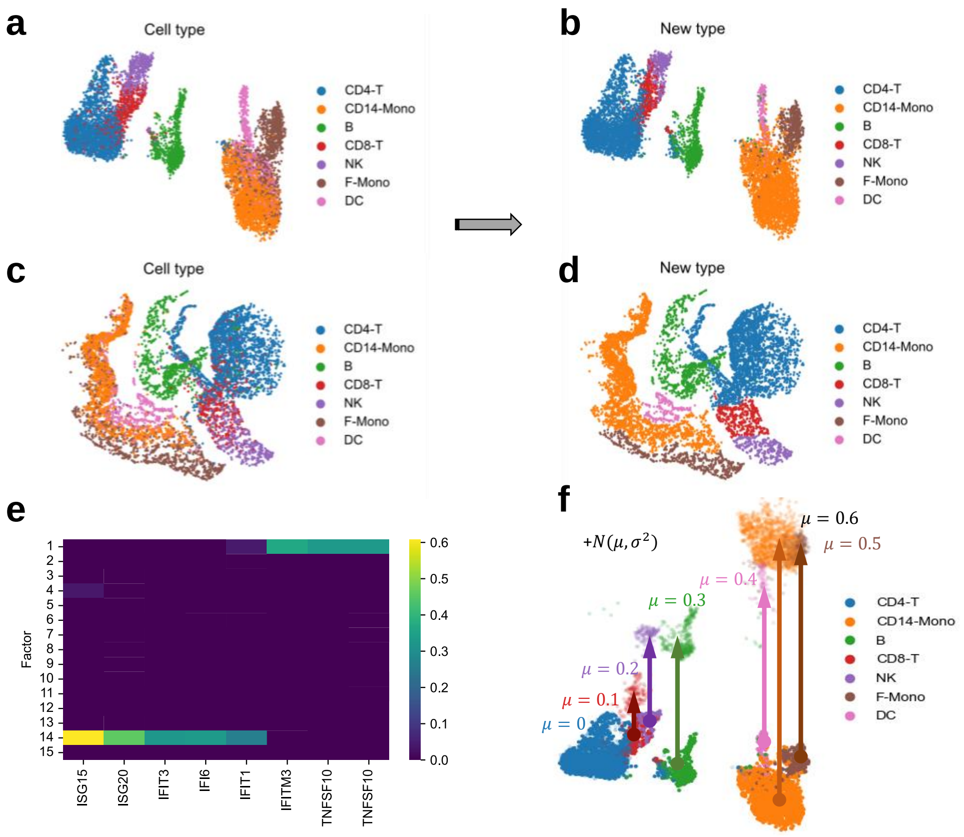

**Supplementary Figure 3. The simulation setup in the PBMC dataset.**

(a, c) UMAP visualizations of the original data (a) and the basal state (c) of control cells, colored by cell types. In the simulation, we aim to test the accuracy of perturbation effect estimation and inference by adding different perturbation effects on seven cell types. Therefore, the cell type labels were adjusted manually to make the cell types have better separation.

(b, d) Similar plots with (a,c) colored by adjusted cell type labels.

(e) The factor loadings of IFN-β associated genes.

(f) The sketch map of the simulation setup. We randomly selected 50% control cells as pseudo-stimulated cells, and added perturbation effects from 0 to 0.6 to their values of factor 14 expression ($z'$) based on their cell type labels and a Gaussian noise with $\sigma = 0.01$.

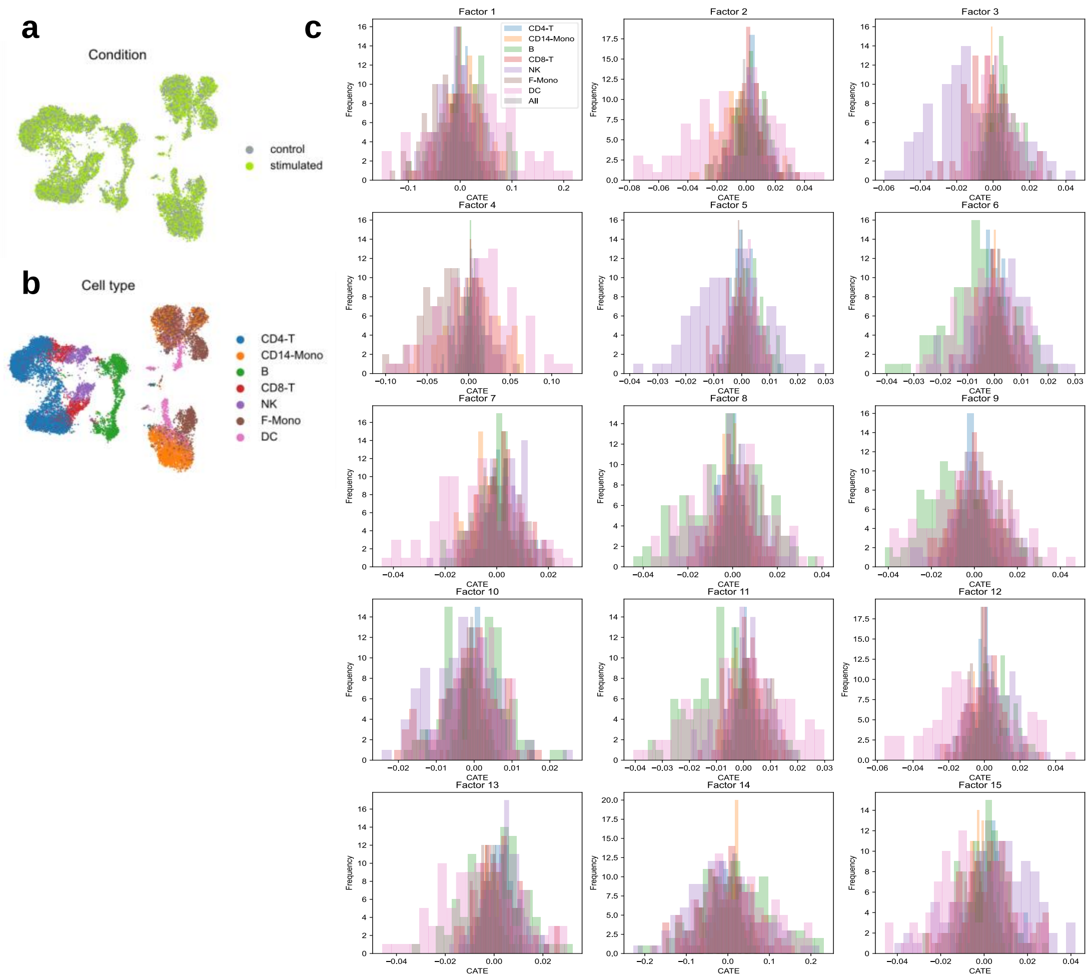

**Supplementary Figure 4. Result of scCAPE estimation in the null (zero perturbation effects) simulation based on the PBMC dataset.**

(a, b) UMAP visualizations of the simulated data, colored by conditions (top) and cell types (bottom). The condition labels of all cells are randomly permuted. We generated 100 random datasets for scCAPE analysis.

(c) The histograms of scCAPE perturbation effect estimations on each factor, averaged on different cell types and all cells.

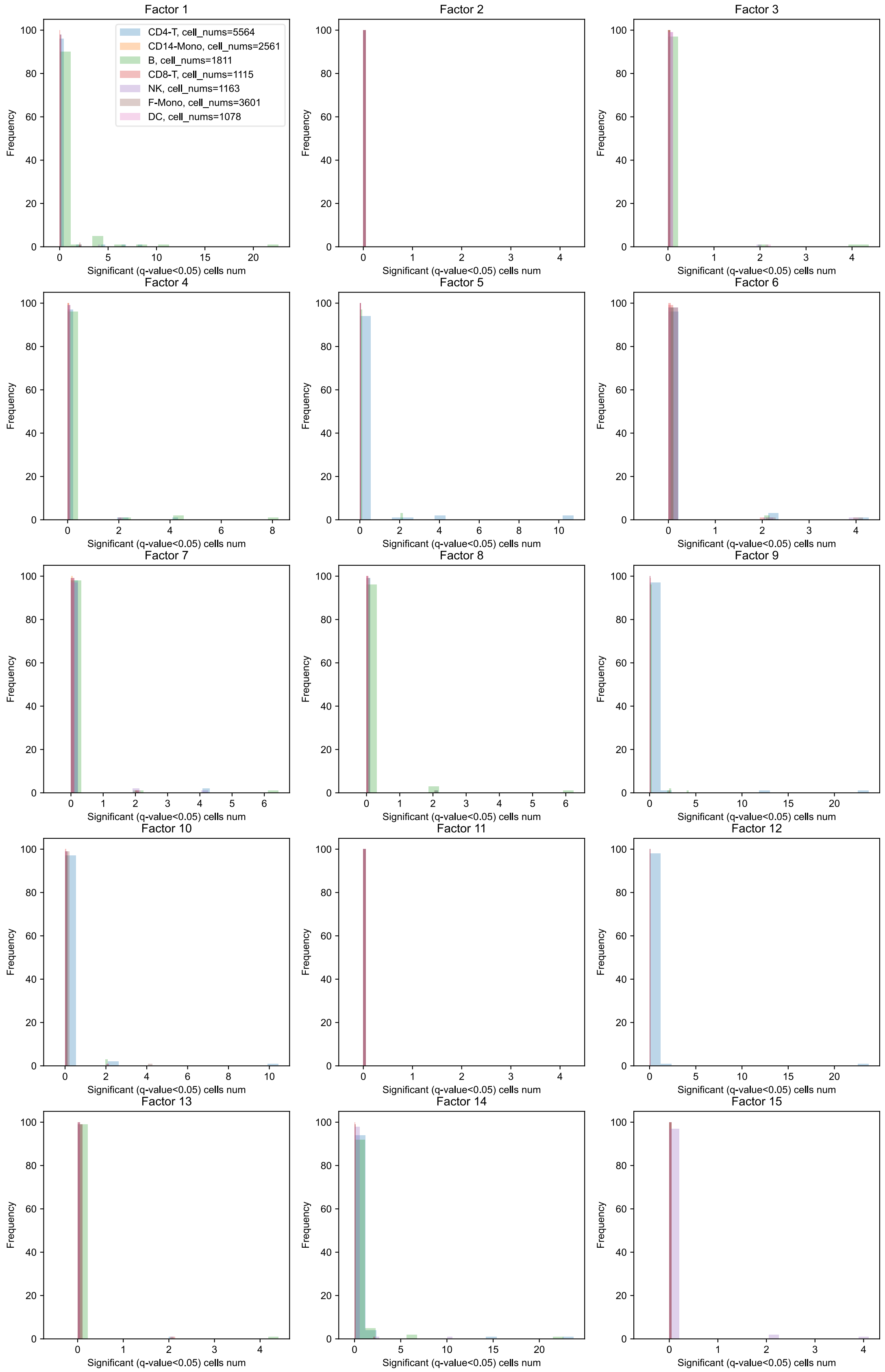

**Supplementary Figure 5. False positive analysis of scCAPE inference on PBMC dataset.** The histograms of the number of significant cells (q-value<0.05) on each factor, colored by different cell types.

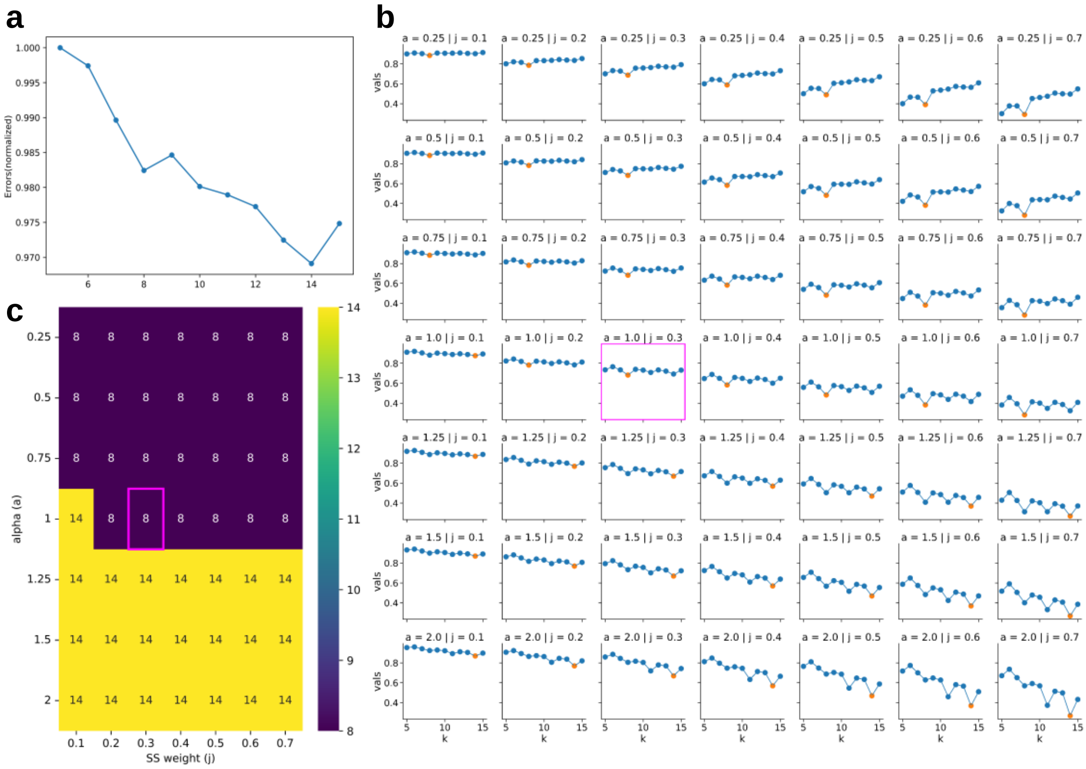

**Supplementary Figure 6. Selection of the number of factors in the T cell CROP-seq dataset.**

(a) Normalized reconstruction errors across different factor numbers.

(b) The value of $f(k;\alpha,j)$ for a varying range of hyperparameter $\alpha$ (rows) and the weight of specificity score $j$ (columns). The highlighted points represent the minimum values. The bounding box shows the curve under the default setting ($\alpha = 1, j = 0.3$).

(c) The optimal number of factors under varying $\alpha$ and $j$. The bounding box shows 8 is the optimal number under the default setting ($\alpha = 1, j = 0.3$).

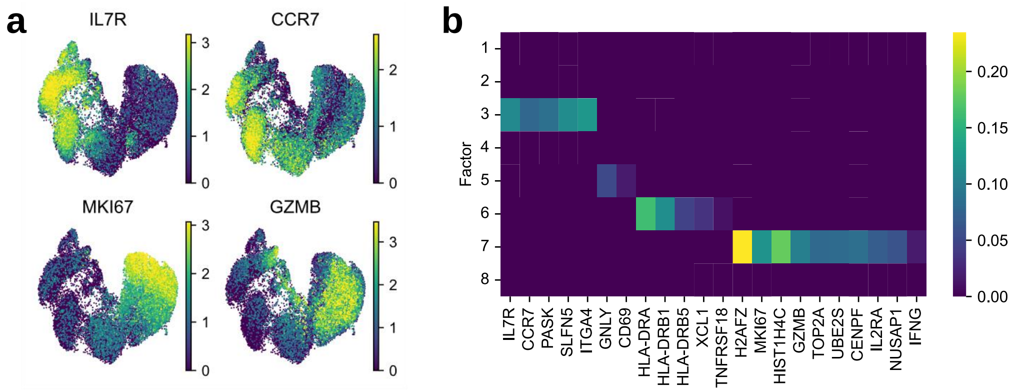

**Supplementary Figure 7. Visualizations and factor annotations of the T cell CROP-seq dataset.**

(a) The expression of unstimulated state markers (IL7R, CCR7) and TCR-stimulated markers (MKI67, GZMB).

(b) The factor loadings of marker genes of T cell resting (IL7R, CCR7, PASK, etc.), activation (CD69, HLA, etc.) and proliferation (H2AFZ, TOP2A, UBE2S, etc.).

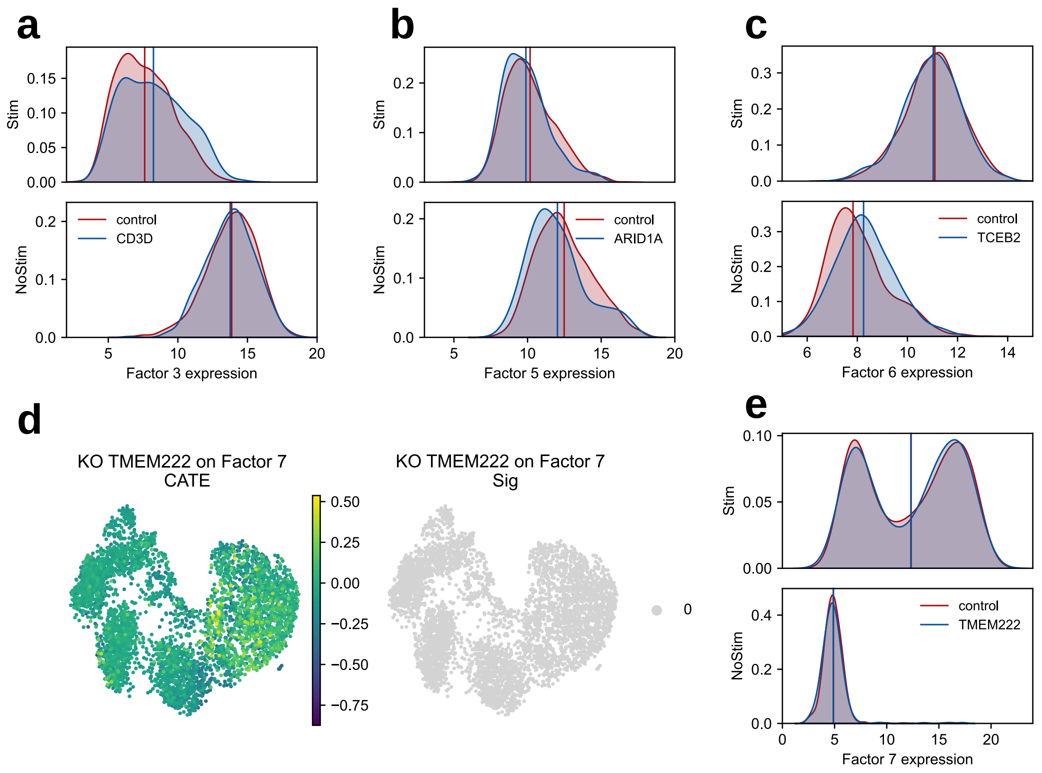

**Supplementary Figure 8. Distinct heterogeneity patterns in perturbation effects between stimulated and unstimulated cells.**

(a) The values of factor 3 in control cells and KO CD3D cells across stimulated and unstimulated conditions, related to Figure 3d;

(b) The values of factor 5 in control cells and KO ARID1A cells across stimulated and unstimulated conditions, related to Figure 3f;

(c) The values of factor 6 in control cells and KO TCEB2 cells across stimulated and unstimulated conditions, related to Figure 3e;

(d, e) The perturbation effects of KO TMEM222 on factor 7. The estimation of perturbation effect and the statistical significance is shown in (d), and the contrast of values of factor 7 in stimulated and unstimulated cells is shown in (e).

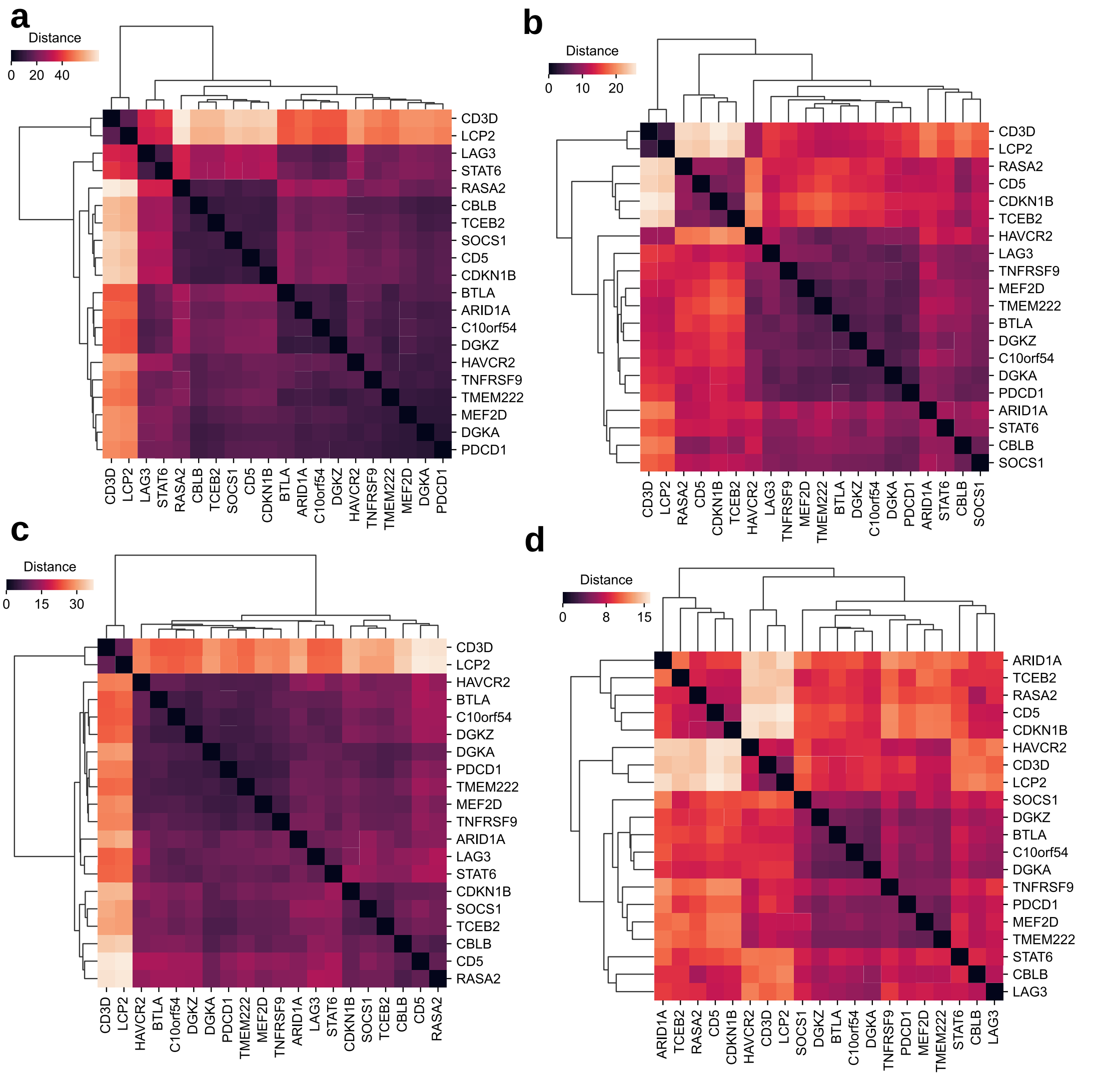

**Supplementary Figure 9. Heatmaps of similarity of different perturbations in the T cell CROP-seq dataset.**

(a, b) Perturbation similarities of factor 7 (T cell proliferation), aggregated over stimulated cells (a) and unstimulated cells (b);

(c, d) Overall perturbation similarities, aggregated over stimulated cells (c) and unstimulated cells (d).

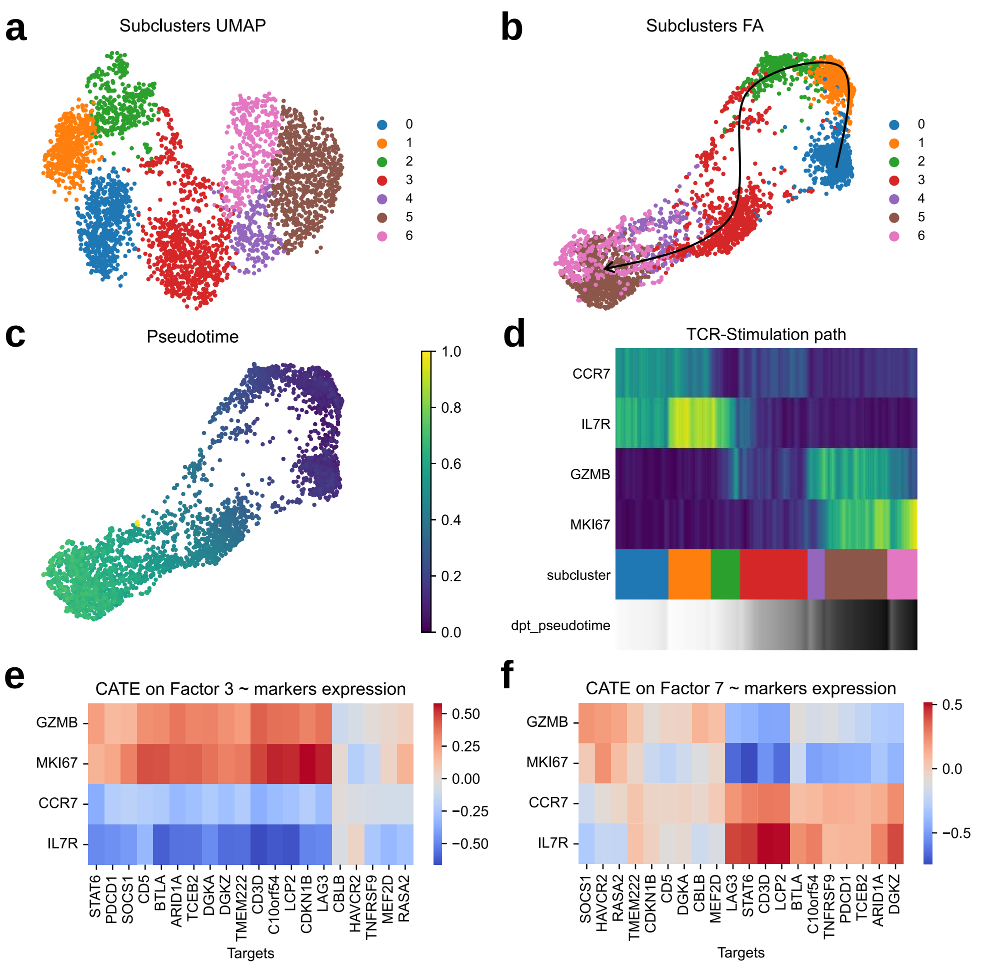

**Supplementary Figure 10. Stimulation trajectory results in the T cell CROP-seq dataset.**

(a) UMAP visualizations of the original data, colored by subclusters using Leiden algorithm.

(b) Force-directed graph drawing (FA, ForceAtlas2[14]) visualizations of the original data, colored by subclusters. The plot is obtained by using '*scanpy.tl.draw_graph*'.

(c) Force-directed graph drawing (FA, ForceAtlas2) visualizations of the original data, colored by pseudotime. The diffusion pseudotime is inferred by using '*scanpy.tl.dpt*'.

(d) The expression of unstimulated state markers (IL7R, CCR7) and TCR-stimulated markers (MKI67, GZMB) along the TCR-stimulation trajectory.

(e) The correlation between the perturbation effects of all targets on factor 3 (T cell resting) and the expression of four marker genes (GZMB, MKI67, CCR7 and IL7R).

(f) The correlation between the perturbation effects of all targets on factor 7 (T cell proliferation) and the expression of four marker genes (GZMB, MKI67, CCR7 and IL7R).

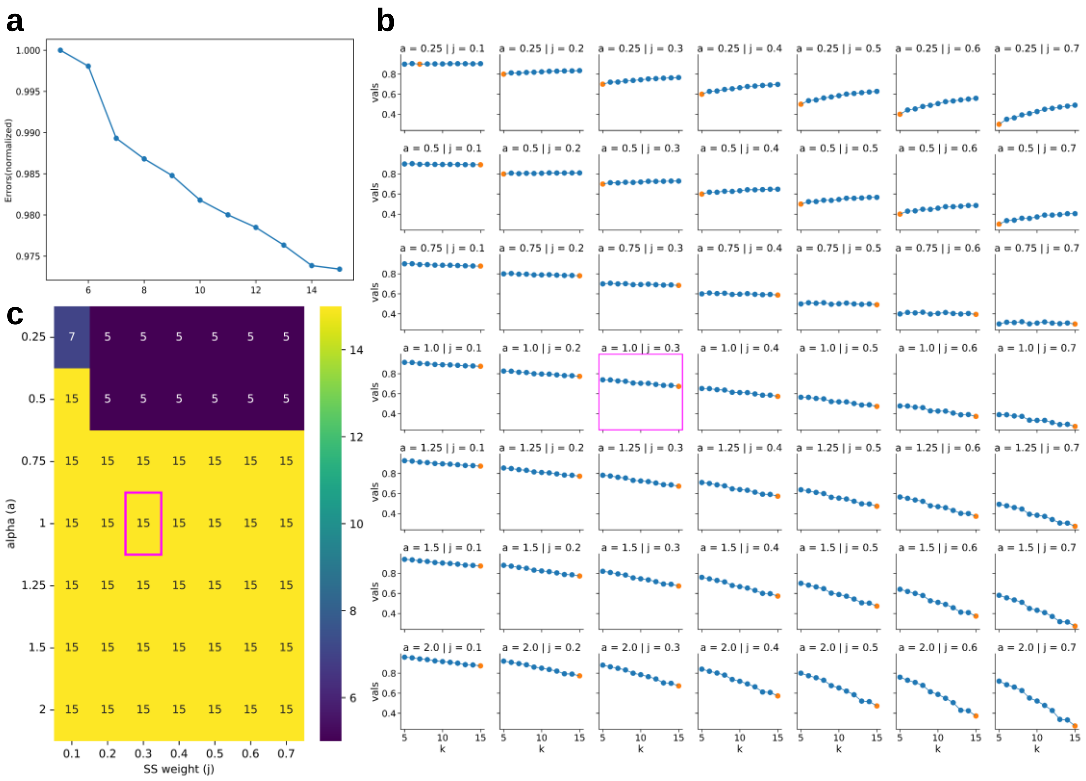

**Supplementary Figure 11. Selection of the number of factors in the K562 Perturb-seq dataset.**

(a) Normalized reconstruction errors across different factor numbers.

(b) The value of $f(k; \alpha, j)$ for a varying range of hyperparameter $\alpha$ (rows) and the weight of specificity score $j$ (columns). The highlighted points represent the minimum values. The bounding box shows the curve under the default setting ($\alpha = 1, j = 0.3$).

(c) The optimal number of factors under varying $\alpha$ and $j$. The bounding box shows 15 is the optimal number under the default setting ($\alpha = 1, j = 0.3$).

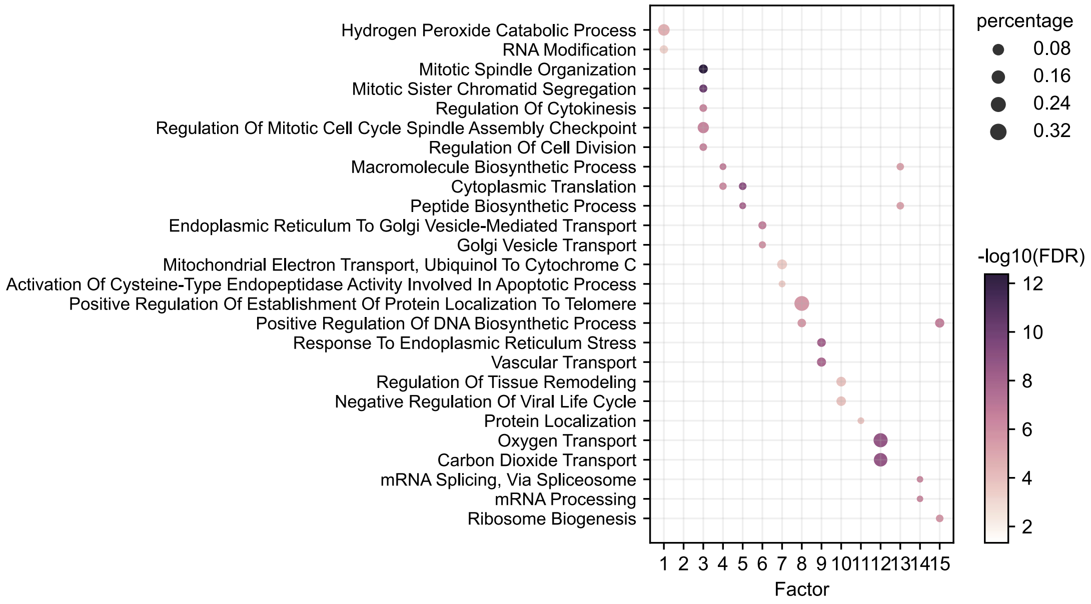

**Supplementary Figure 12. Factor annotations in the K562 cell Perturb-seq dataset.** Factor function annotations based on gene set enrichment analysis. The size of the dot represents the percentage of selected genes in that pathway or gene ontology term. The color of the dot shows the -log10(adjusted p value).

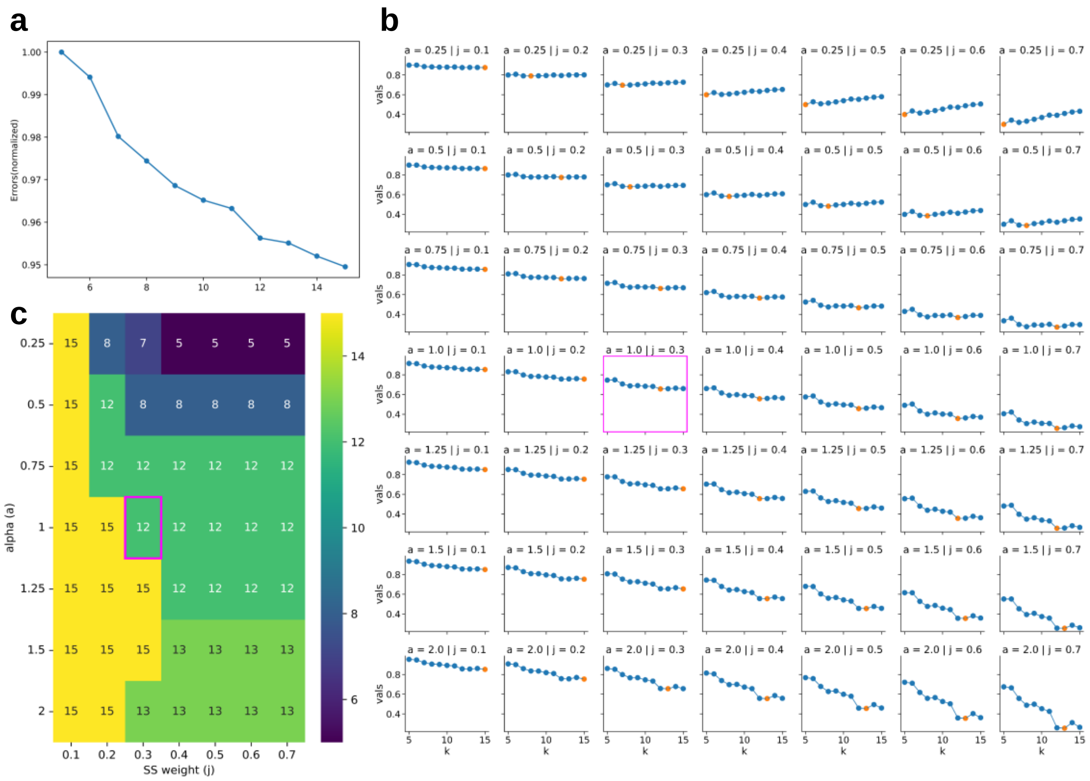

**Supplementary Figure 13. Selection of the number of factors in the high-MOI dataset.**

(a) Normalized reconstruction errors across different factor numbers.

(b) The value of $f(k;\alpha,j)$ for a varying range of hyperparameter $\alpha$ (rows) and the weight of specificity score $j$ (columns). The highlighted points represent the minimum values. The bounding box shows the curve under the default setting ($\alpha = 1, j = 0.3$).

(c) The optimal number of factors under varying $\alpha$ and $j$. The bounding box shows 12 is the optimal number under the default setting ($\alpha = 1, j = 0.3$).

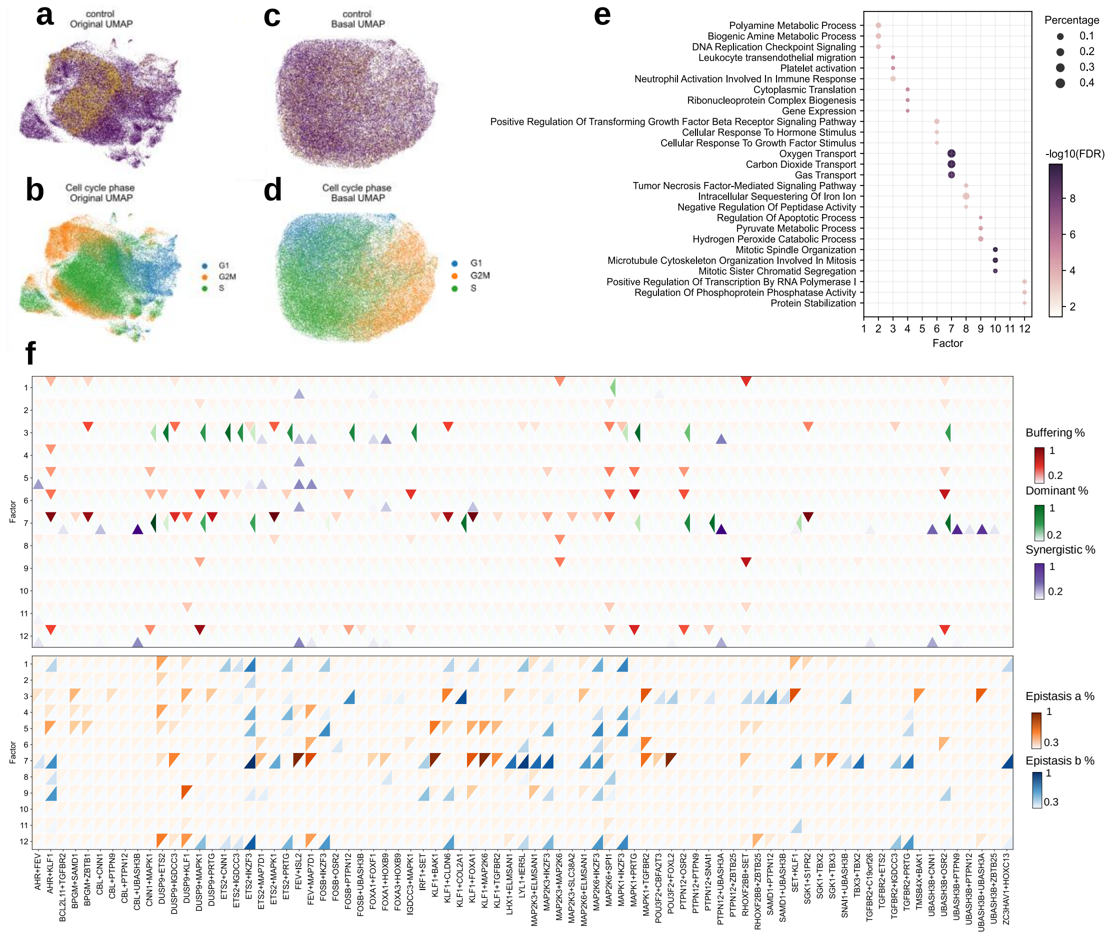

**Supplementary Figure 14. Additional results in the high-MOI Perturb-seq dataset.**

(a) UMAP visualizations of the original data, colored by control indicators (yellow: control cells, purple: perturbed cells).

(b) UMAP visualizations of the original data, colored by cell cycle phases. The cell cycle phases are inferred using ‘scanpy.tl.score_genes_cell_cycle’.

(c, d) UMAP visualizations of scCAPE basal state, colored by control indicators (c) (yellow: control cells, purple: perturbed cells) and cell cycle phases (d). The basal state removes perturbation information and retains the local cell state variations (cell cycle phases).

(e) Annotations of factor function based on gene set enrichment analysis. The size of the dot represents the percentage of selected genes in that pathway or gene ontology term. The color of the dot shows the -log10(adjusted p-value).

(f) Inferences of factor-specific genetic interactions.

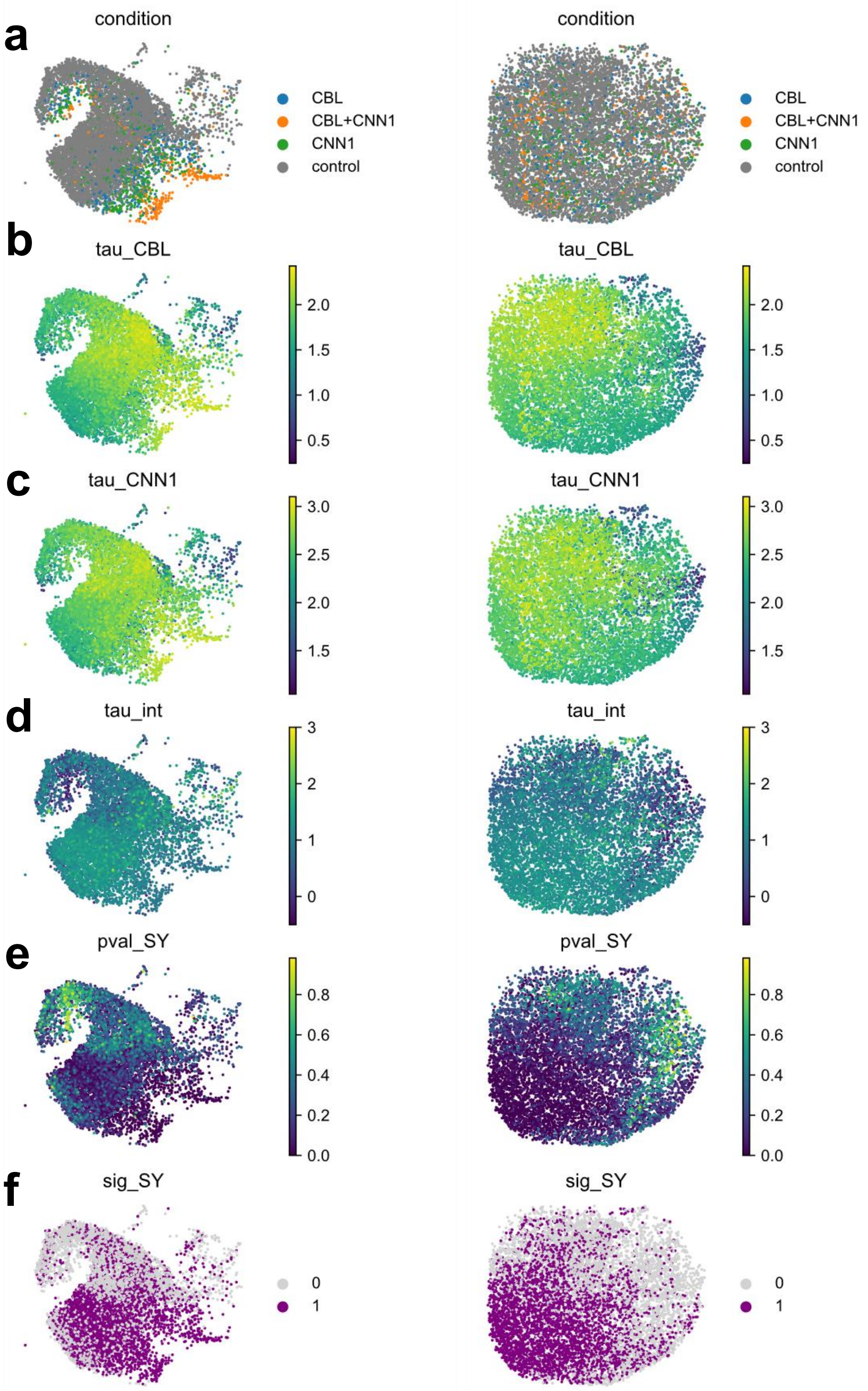

**Supplementary Figure 15. UMAP visualizations of the synergistic interactions between CBL and CNN1 in driving erythroid differentiation.**

The left and right panel are UMAP visualizations of the original data and the basal state, colored by perturbation labels (a), the perturbation effect of CBL ($\hat{\tau}_a$) (b), the perturbation effect of CNN1 ($\hat{\tau}_b$) (c), estimation of interaction term ($\hat{\tau}_{int}$) (d), adjusted p-values of the synergistic test (e), and the significance of the synergistic test under $\alpha = 0.05$ (f).

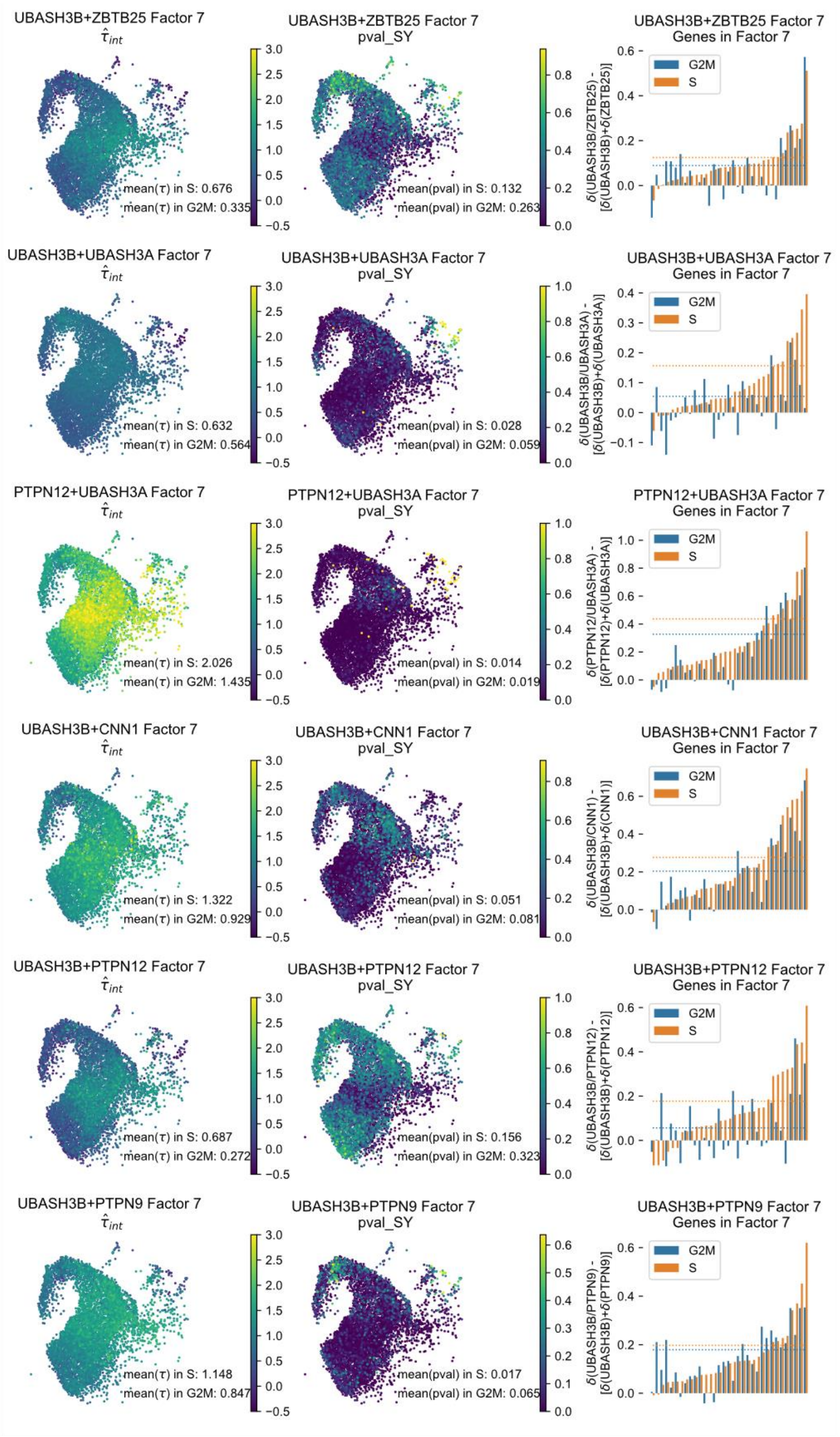

**Supplementary Figure 16. Synergistic test results for the other erythroid drivers.**
UMAP visualizations of the original data, colored by the estimation of the interaction term ($\hat{\tau}_{int}$) (left) and adjusted p-values of the synergistic test (middle). The right panel shows the interaction effects of high loading genes in factor 7 averaged on S phase cells (orange) and G2M phase cells (blue). The dotted line denotes the average interaction value across all 33 high loading genes in factor 7. $\delta(\cdot)$ denotes average deviation in gene expression relative to control cells, and $\delta(a/b) - [\delta(a) + \delta(b)]$ represents average interaction effects on these high-loading genes.

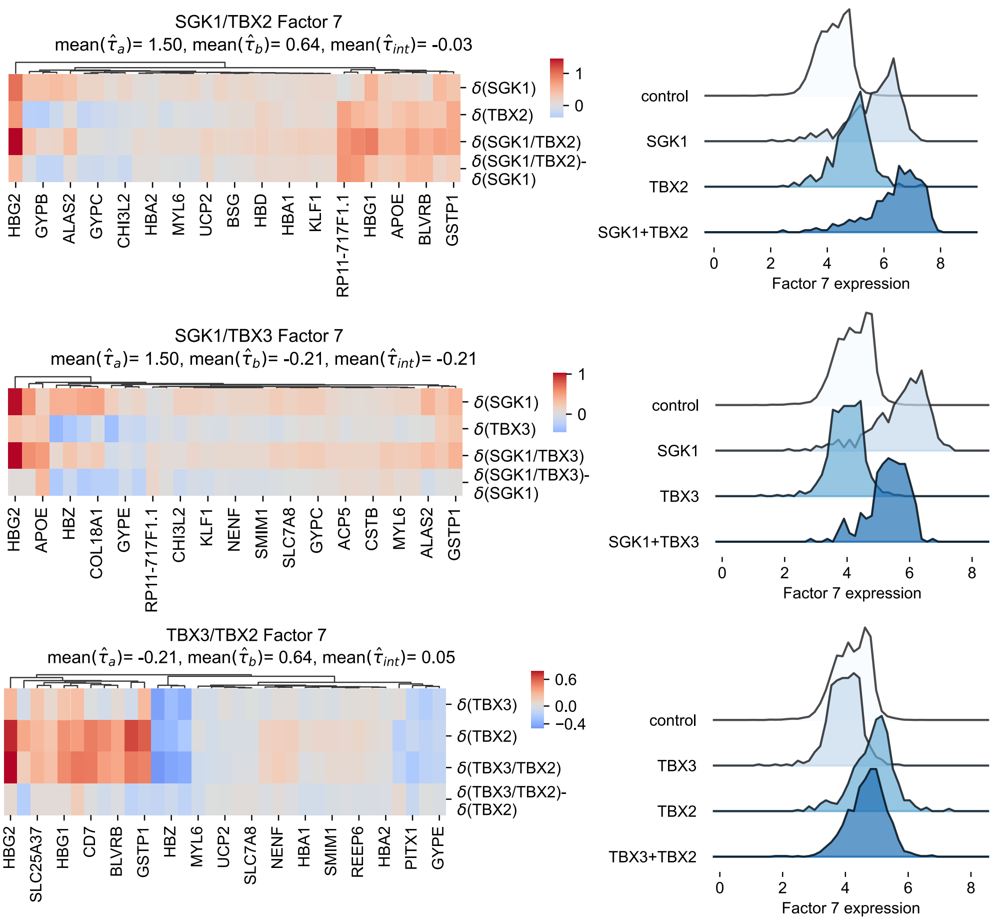

**Supplementary Figure 17. Epistasis interactions among SGK1, TBX2 and TBX3.**
We compared values of factor 7 of four conditions (right) and the expression of high-loading genes in factor 7 (left). $\delta(\cdot)$ denotes average deviation in gene expression relative to control cells.

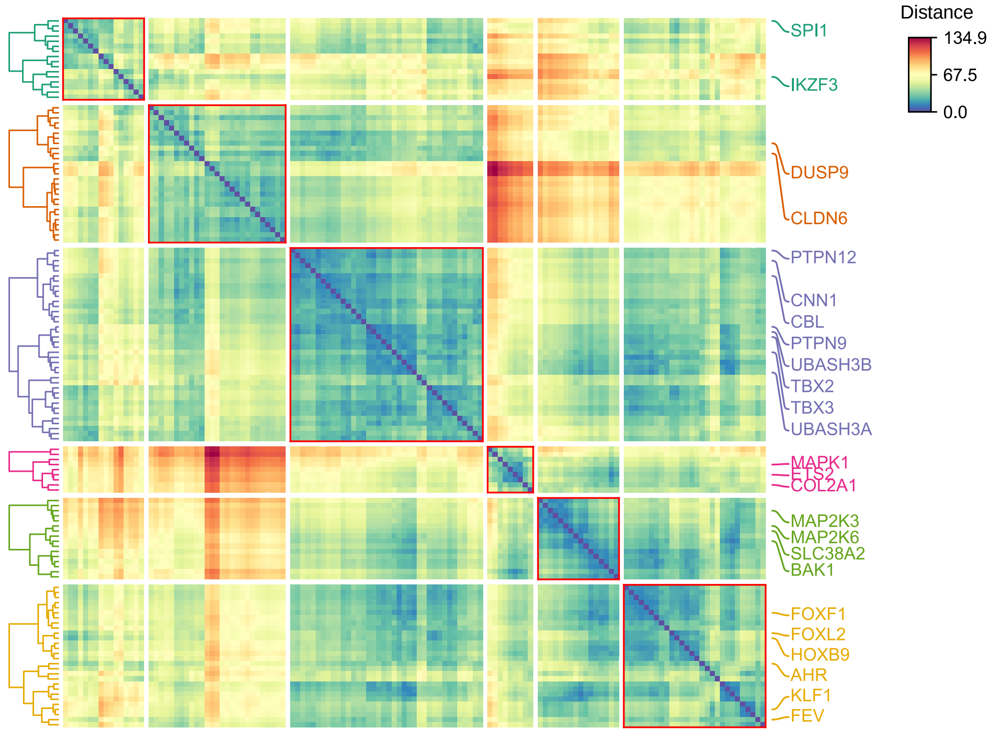

**Supplementary Figure 18. The overall similarity between perturbations in the high-MOI dataset, aggregated over all cells.**

Erythroid differentiation drivers (such as CNN1, CBL, UBASH3A/B), megakaryocyte differentiation drivers (such as MAPK1 and ETS2) and some pioneer factors (such as HOXB9, FOXF1 and FOXL2) are distinctly delineated.

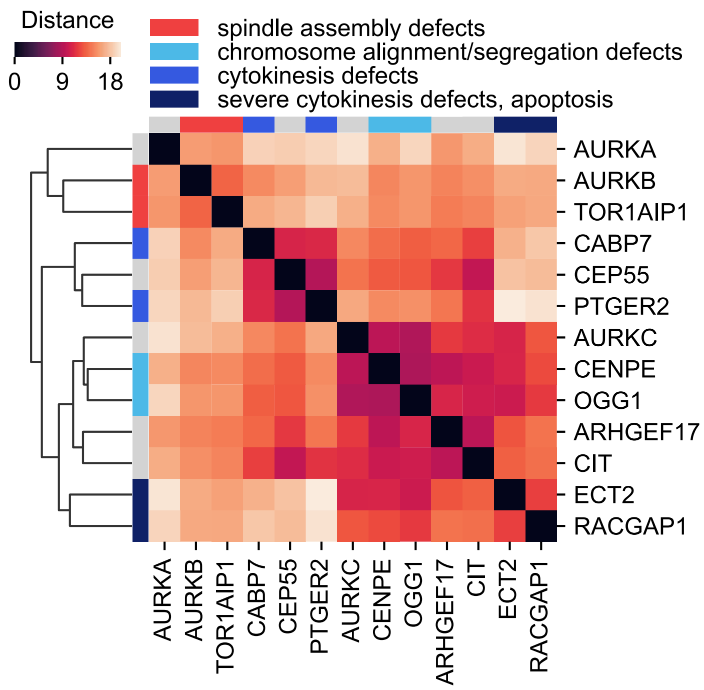

**Supplementary Figure 19. The overall similarity between perturbations in Perturb-seq K562 cell dataset, aggregated over all cells.**

The categorization of perturbations based on their overall regulatory effects predominantly aligns with their groupings by cellular morphology in HeLa cells[15]. The two primary clusters correspond to early mitotic defects with errors in spindle assembly (AURKB and TOR1AIP1) and chromosome segregation abnormalities or cytokinesis failures with binuclear or polylobed phenotypes (such as CIT, PTGER2, RACGAP1) respectively. Within the second cluster, CENPE and OGG1 exhibit chromosome alignment and/or segregation problems, while CABP7 and PTGER2 display normal chromosome segregation but appear to be specifically affected in cytokinesis.

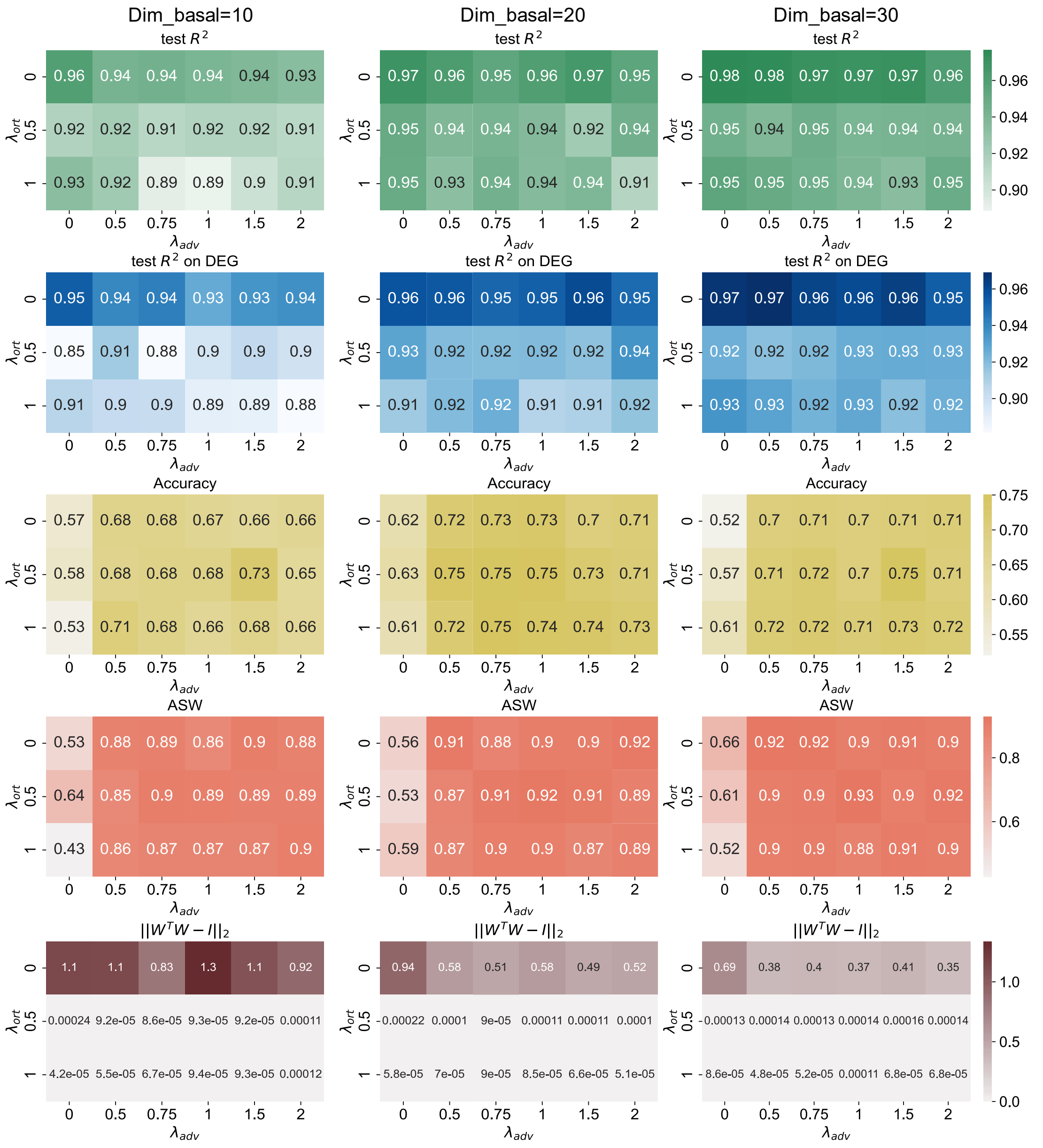

**Supplementary Figure 20. The influences of varying hyperparameters on reconstruction and disentangling performance.**

In the PBMC dataset, we evaluated the influence of varying hyperparameters on the performance of scCAPE, including the dimensions of the basal state, the coefficient of the adversarial loss ($\lambda_{adv}$) and the coefficient of the orthogonal constraint ($\lambda_{ort}$). The metrics include (1) test $R^2$, the squared Pearson correlation between the reconstruction and the real mean expression averaged on cell states clustered using the Leiden algorithm for the test dataset; (2) test $R^2$ on DEG, the mean squared Pearson correlation between the reconstruction and the real expression of the top 50 DEGs for the test dataset; (3) Accuracy, the percentage of stimulated cell which has the same cell type label with its nearest control neighbor in the basal state; (4) ASW, the average silhouette width, where higher values indicate better mixing performance; (5) The orthogonality of the loading matrix. Under each hyperparameter setting, we trained the networks for 5 times and reported the average results.
